# Testing the evolutionary potential of an alpine plant: Phenotypic plasticity in response to growth temperature far outweighs parental environmental effects and other genetic causes of variation

**DOI:** 10.1101/2024.02.20.581287

**Authors:** Pieter A. Arnold, Shuo Wang, Rocco F. Notarnicola, Adrienne B. Nicotra, Loeske E. B. Kruuk

## Abstract

Phenotypic plasticity and rapid evolution are fundamental processes by which organisms can maintain their function and fitness in the face of environmental changes. Here we quantified the plasticity and evolutionary potential of an alpine herb *Wahlenbergia ceracea*. Utilising its mixed-mating system, we generated outcrossed and self-pollinated families that were grown in either cool or warm environments, and that had parents that had also been grown in either cool or warm environments. We then analysed the contribution of environmental and genetic factors to variation in a range of phenotypic traits including phenology, leaf mass per area, photosynthetic function, thermal tolerance, and reproductive fitness. The strongest effect was that of current growth temperature, indicating strong phenotypic plasticity. All traits except thermal tolerance were plastic, whereby warm-grown plants flowered earlier, grew larger, produced more reproductive stems compared to cool-grown plants. Flowering onset and biomass were heritable and under selection, with early flowering and larger plants having higher relative fitness. There was little evidence for transgenerational plasticity, maternal effects, or genotype-by-environment interactions. Inbreeding delayed flowering and reduced reproductive fitness and biomass. Overall, we found that *W. ceracea* has the capacity to respond rapidly to climate warming via plasticity, and the potential for evolutionary change.

**Highlight:** We found strong plasticity to growth environment in many phenotypic traits, but little effect of parental environment, revealing capacity to respond rapidly to climate warming, and potential for evolutionary change.

## Introduction

Climate change is exposing organisms to ever increasing mean temperatures and more frequent extreme events (Harris *et al*., 2018). Temperature is a universal, pervasive environmental variable that limits a species’ occupiable niche (Walther, 2003; Tattersall *et al*., 2012; Nievola *et al*., 2017). It is therefore unsurprising that steadily warming mean temperatures have been observed to affect biological processes, such as phenology, globally (Walther *et al*., 2002; Parmesan and Yohe, 2003). For example, in warmer years the date of flowering onset in plants occurs considerably earlier, and has been shown to be tightly related to the pattern of mean climatic warming (Parmesan and Hanley, 2015), especially in high elevation plants (Giménez-Benavides *et al*., 2011; Dorji *et al*., 2020). Indirect reproductive effects due to phenological changes from warming, as well as direct effects on fertility and reproductive traits, can also have consequences for individual- and population-level fitness (Anderson, 2016). To determine the realised impact of warming on natural selection and evolutionary processes, we must assess how warming affects phenotypic responses and fitness, and how these vary among individuals in a population.

There are several mechanisms, not mutually exclusive, by which plants can tolerate climatic and environmental changes. Plants could already have a high natural resilience to warmer temperatures, thereby partially or completely avoiding thermal stress (De Kort *et al*., 2020; Andrew *et al*., 2023). Alternatively, phenotypic plasticity – the capacity of a single genotype to exhibit multiple phenotypes based on changes in the environment (Bradshaw, 1965, 2006) – could facilitate short-term changes to their phenotype, to limit exposure to stress effects or to mitigate damage (Nicotra *et al*., 2010; Fox *et al*., 2019; Brooker *et al*., 2022). For instance, exposure to increased mean temperatures can benefit some plants by acclimating (or priming) them to face more stressful conditions in a more prepared state, so that extreme events have a lesser impact (Hilker *et al*., 2016). Plants generally have limited capacity to move to escape unfavourable temperatures due to the constraints and costs of dispersal, therefore responding to local conditions through plasticity is likely to be evolutionarily favoured (Edelaar *et al*., 2017).

The widespread occurrence of phenotypic plasticity generates assumptions that it is an evolutionary adaptation to environmental heterogeneity (Hendry, 2016). In other words, selection will favour plasticity because it improves an individual’s performance or fitness in a particular environment and would therefore be considered adaptive (Bonser, 2021). Here, Genotype × Environment interactions (G×E) specifically refers to variation in plastic responses among genotypes within a population. That is, the expression of phenotypes in response to the environment (E) varies depending on the genotype (G). Although phenotypic plasticity is at times seen as equivalent to G×E, plasticity can result simply in response to variation in E, without there being significant variation attributable to G. For plasticity to evolve, however, there has to be genetic variation in plasticity on which selection can act, hence a significant G×E interaction is a requisite for demonstrating the adaptive significance of plasticity (Josephs, 2018). Yet, evidence suggests that plasticity being costly is about as common as it is being beneficial; the exact balance will depends on the severity of the stress and the relative strengths of the costs and benefits of plasticity (van Buskirk and Steiner, 2009; Auld *et al*., 2010; Hendry, 2016). Indeed, there is emerging evidence that trait canalisation (i.e., no plasticity, also called robustness) in response to warming may be favoured when costs of plasticity are high or plasticity is maladaptive (Stinchcombe *et al*., 2004; Arnold *et al*., 2019b; Svensson *et al*., 2020).

While plasticity in response to warming is common in plants (Nicotra *et al*., 2010), the degree to which different traits respond can differ markedly. From a global meta-analysis, plasticity in leaf morphology, plant biomass, and several physiological traits (including chlorophyll content and photosynthetic efficiency: *F*_V_/*F*_M_ and φPSII) all indicate strong responses to increasing mean annual temperature (Stotz *et al*., 2021). However, within a species there may be substantial inter-trait variation in responses. For example, the alpine herb *Wahlenbergia ceracea* exhibits different patterns of plasticity in response to temperature, depending on trait type. Warmer, but not stressful, growth temperatures increase leaf mass per area (LMA), heat tolerance traits, and reproductive output (flowers and capsules), and also induce earlier flowering, decreased photosynthetic efficiency, and lowered biomass (Arnold *et al*., 2022). Along with the variation in plastic responses among traits of different types, there is also substantial intraspecific variation in plastic responses to temperature (Arnold *et al*., 2022).

The patterns of selection acting on plant traits may also be affected by climate and environmental factors. Climate change has already had widespread effects on plant traits, especially the timing of reproductive events, where earlier flowering under warmer conditions almost ubiquitously increases fitness and is therefore under strong directional selection (Franks *et al*., 2007; Anderson *et al*., 2012; Anderson, 2016; Wadgymar *et al*., 2018b; Ehrlén and Valdés, 2020). The relationship between functional traits and fitness depends on the environment, yet other than phenology, empirical tests of selection on functional traits due to climate warming are uncommon (Geber and Griffen, 2003; Kimball *et al*., 2012). Totland (1999) found that *Ranunculus acris* were under selection for more flowers in both control and warmed field conditions, but selection for larger leaf size was only apparent in control conditions. Using an urban (warmer and drier) environment as a climate change analogue to test for its effect on selection, Lambrecht *et al*. (2016) found strong evidence for selection on functional traits, including increased plant size, leaf number, specific leaf area, and later senescence in urban conditions in *Crepis sancta*. Establishing which phenotypic traits are correlated with fitness (i.e., under selection) under benign and stressful conditions is a crucial component for understanding the potential for rapid evolutionary responses (Anderson, 2016). For evolutionary change to keep pace with climate change, the phenotypic trait under selection must also have heritable variation on which selection can act (Scheiner *et al*., 2020). However, there are also alternative mechanisms of affecting phenotype change across generations.

Transgenerational plasticity or parental environmental effects can be demonstrated when the conditions to which a parent is exposed shape offspring phenotype function and fitness in their environment (Mousseau and Fox, 1998; Bonduriansky, 2021), most cases of which are mediated by maternal effects (Herman and Sultan, 2011). A factorial design of at least two offspring and two parental environments that match or mismatch allows transgenerational and within-generation plasticity to be tested simultaneously (Uller *et al*., 2013). Adaptive transgenerational plasticity (also called ‘anticipatory parental effects’) theory posits that selection on parental responses to their environment confers benefits to the offspring when their environment matches that of the parents, particularly when the environmental conditions are stressful (Uller, 2008; Herman and Sultan, 2011). Conversely, when the parent and offspring environments are not stable or are unpredictable, or when they mismatch (e.g., due to change in season or annual change), there may be a cost to the offspring of producing a phenotype that reflects the parental environment rather than the current environment (Engqvist and Reinhold, 2016).

A previous meta-analysis has revealed little evidence for transgenerational plasticity conferring a clear benefit in plants, especially when close proxies for fitness are used (Uller *et al*., 2013). Yet, there are cases of strong parental effects in response to temperature in plants. For example, Whittle *et al*. (2009) found *Arabidopsis thaliana* plants substantially increased reproductive output under relatively hot conditions (30°C) when prior generations had also been grown in the same hot conditions. There is also generally stronger evidence for transgenerational plasticity affecting early life traits like seed germination. For example, Wadgymar *et al*. (2018a) showed greater and more variable transgenerational plasticity than within-generation plasticity in germination of *Boechera stricta* plants across an elevation gradient. The effects of parental temperature in *W. ceracea* have also been found to affect germination and dormancy patterns, but to a lesser extent than the temperatures in which seeds germinated (Wang *et al*., 2021; Notarnicola *et al*., 2023b).

Climatic warming is expected to increase rates of self-pollination in plants with mixed-mating strategies (i.e., those that can facultatively self-pollinate in the absence of cross-pollination), due to phenological mismatches with pollinators (Hegland *et al*., 2009). However, while self-pollination is a contingency strategy that may assure reproductive success in the face of climate warming, inbreeding depression may be worsened in stressful environments (Armbruster and Reed, 2005) and it also reduces adaptive potential (Hamann *et al*., 2021). While the effects of inbreeding under some stressors associated with climate change (drought, herbivory, nutrient deficiency) have been studied in mixed-mating species (Hamann *et al*., 2021), to our knowledge only one study has investigated the effects of inbreeding with warming in a mixed-mating species (Wang *et al*., 2021).

In the current study, we used a large-scale glasshouse experiment to measure a suite of phenotypic traits on outcrossed and self-pollinated *W. ceracea* plants, from families that were grown in either cool or warm environments that had parents that were grown in either cool or warm environments. We addressed the following questions (Q):

1. To what extent do a suite of traits respond to growth temperature through phenotypic plasticity?
2. Is there any evidence of transgenerational plasticity or benefits for offspring that are grown under conditions that match the conditions in which their parents were grown?
3. Are the phenotypic traits heritable, and are there either maternal effects or G×E interactions?
4. What is the direction and strength of selection on the traits and how does it vary with growth temperature?
5. Are there effects of inbreeding on the suite of traits, and does any inbreeding depression vary with temperature?

For Q1, we hypothesised that the warmer growth temperature would be stressful, reducing plant function and biomass, but that plants would also respond by flowering earlier, which may compensate to result in equal or higher fitness, and that acclimatory processes would improve heat tolerance. For Q2, we hypothesised that the parental environment would have a small effect on offspring phenotype (i.e., development of seed under parental temperatures that matched the offspring growth temperature would benefit fitness compared to mismatched offspring). For Q3, we hypothesised that the heritable traits would be flowering onset, biomass, LMA, and heat tolerance, based on previous findings of highly plastic responses to temperature and intraspecific variation in these responses (Arnold *et al*., 2022). For Q4, we hypothesised that traits under selection would be the same as those heritable in Q3. For Q5, we predicted that there would be evidence for inbreeding depression primarily in fitness. Addressing this series of questions together will elucidate the evolutionary potential of functional responses to warmer growth temperatures that are expected under future climate change.

## Materials and methods

### Species description, seed source, and F0-F2 generations

*Wahlenbergia ceracea* Lothian (Campanulaceae) waxy bluebell is a short-lived, protandrous, and facultatively autogamous biennial herb that is sparsely distributed across south-eastern Australia and Tasmania (Nicotra *et al*., 2015). For this study, seeds were collected between 1590 m and 2100 m a.s.l. from Kosciuszko National Park, NSW, Australia (36.43°S, 148.33°E) in 2015 and 2016 (see Notarnicola *et al*., 2021 for further details), and brought to The Australian National University, Canberra, ACT, Australia. These seeds formed the F0 generation: we describe below their rearing and subsequent breeding design for F1, F2, and F3 generations, with the F3 plants then being used for the analyses here of phenotypic plasticity, transgenerational plasticity, heritable genetic variance, and inbreeding depression.

### F0, F1, and F2 generations

Detailed descriptions of the conditions and breeding design for producing plants for generations F0 to F3 in this study have been given previously (Notarnicola *et al*., 2021; Wang *et al*., 2021; Arnold *et al*., 2022). Briefly, F0 plants grown from field-collected seeds were raised in glasshouses that mimicked average alpine summer temperatures (25/15°C during germination and growth, which was reduced to 20/15°C at peak flowering). F1 plants were produced by crossing 48 F0 plants as pollen donors with 48 F0 plants as ovule donors (96 plants from 63 unique F0 families) to produce 48 F1 families by hand-pollination. F1 plants were raised under the same conditions as the F0 plants (Supplementary Fig. S1). The F2 generation was produced using a partial-diallel (maximising the number of parents used to generate families) half-sib breeding design in which 12 F1 plants were used as pollen donors and each outcrossed with at least four unrelated pollen receivers (48 pollen receivers in total). F2 plants were grown in glasshouses under two temperature regimes (‘*cool*’: 20/15°C day/night conditions and ‘*warm*’ 30/25°C day/night conditions) that hereafter constitute the ‘*parental temperature*’. These two temperature regimes represent the cool conditions to which these alpine plants are already well-adapted, and the warm conditions simulate substantial warming that could occur under an extreme climate change scenario.

### F3 breeding design and pedigree

Within each parental temperature treatment, 12 F2 lines were assigned as pollen donors and these were each outcrossed with four unrelated pollen receivers, such that cool and warm F2 lines were full-sibs of the same F1 parentage. Each of the F2 parent plants in each treatment were also self-pollinated to generate 12 F3 inbred maternal and 11 inbred paternal lines, in addition to the 12 outcrossed F3 lines that were reciprocally outcrossed from paired F2 pollen donors from each parental temperature treatment (Supplementary Fig. S1). In total, there were 96 outcrossed F3 families and 46 inbred F3 families, all controlled by hand-pollination and bagging flowers. The capsules that were formed after hand-pollination of the F2 plants were removed after they had opened, dried, and the seeds had browned. All seeds taken from these capsules were stored in a desiccator for at least seven weeks at ∼20°C and 15-20% relative humidity before the beginning of the experiment.

### F3 growth experiment and temperature treatments

For each F3 cross, 20-30 seeds from a single capsule were sown across two 50 mm Petri dishes containing 1% agar, each corresponding to a growth temperature treatment. The dishes were sealed and moved to an incubator for cold stratification at 5°C in darkness for six weeks to release seeds from dormancy, thereby improving subsequent germination success (Wang *et al*., 2021). At least eight healthy seedlings per dish were transplanted into punnets containing seed raising mix (Debco Pty Ltd, VIC, Australia) and moved into ideal common glasshouse conditions (25/18°C) to grow for two months. The 25/18°C temperature regime was chosen based on the daytime temperature that increased growth rates of *W. ceracea* seedlings (Arnold *et al*., 2022), while maintaining natural cooling at night. Liquid fertiliser (Thrive Soluble All Purpose Plant Food; Yates, NSW, Australia) at a concentration of 0.5 g L^-1^ was added regularly to promote growth and seedlings were watered twice daily. Up to eight healthy seedlings (6-40 mm in diameter) from each family were transplanted into individual plastic pots (125 mm diameter, ∼600 mL) filled with soil suitable for Australian natives combined with 3 g L^-1^ of low phosphorus slow-release fertiliser (Scotts Osmocote Plus Trace Elements: Native Gardens; Evergreen Garden Care Australia, NSW, Australia).

The potted F3 plants were moved to their *growth temperature* treatment seven days after transplantation (commenced 7 November 2019). For the *cool* treatment, plants were placed in a large glasshouse room set to 20/15°C (day/night) under natural photoperiod and for the *warm* treatment, plants were placed in an adjacent glasshouse room set to 30/20°C (day/night). An automatic shade screen was active between 12:00-14:30 and when external temperatures exceeded 30°C for the cool treatment and 33°C for the warm treatment, to prevent excess solar radiation and overheating of the glasshouses, otherwise plants received natural light. Plants were watered daily initially until growing well and then watered to weight to ensure that the soil did not dry out and water was not limiting. As such, the plants in the warm treatment received more frequent watering than did the cool treatment. Pest treatments (VectoBac larvicide for treating fungus gnats and sulphur evaporation for treating powdery mildew) were conducted as required and liquid fertiliser 1 g L^-1^ was applied approximately fortnightly as required for maintaining healthy growth. Within each treatment, we randomised the distribution of one plant from each family to each of four blocks consisting of 29 columns and five rows, which was replicated across both rooms. Not all families had eight healthy plants, but each had at least five plants that were distributed randomly across both treatments. Empty pots were used in place of a missing plant, so that the position layout was preserved across all blocks. In total, there were 1,024 plants (out of an ideal 1,152) at the beginning of the growth experiment (512 in each growth treatment).

### F3 phenotypic trait measurement

We measured a range of phenotypic traits on the F3 plants grown in the two temperature treatments, where chosen traits characterise many important plant functions, which have potential to respond to environmental factors. The date of the first flower produced by every plant was recorded throughout the experiment, which was checked at least every 2-3 days, and this was converted to the day of *flowering onset* since the beginning of the growth temperature treatments. After four weeks in the treatments (9 January 2020), we began phenotyping leaves for thermal tolerance limits and photosynthetic parameters using chlorophyll fluorescence. We used a Pulse Amplitude Modulated (PAM) chlorophyll fluorescence imaging system (MAXI-Imaging-PAM, Heinz Walz GmbH, Effeltrich, Germany) to measure the *photosystem II (PSII) operating efficiency (φPSII)* and the *maximum quantum efficiency of PSII photochemistry (F_V_/F_M_)*. We also measured the heat and cold tolerance limits of leaves by measuring the temperature-dependent change in minimal chlorophyll fluorescence (*T-F*_0_) using controllable thermoelectric Peltier plates (plate: CP-121HT; controller: TC-36-25; TE-Technology, Inc., Michigan, USA) in conjunction with imaging fluorimeters. We determined the critical threshold temperatures at which *F*_0_ increased rapidly to indicate the impairment of PSII, as measures of physiological *heat tolerance* (*T_crit-hot_*) and *cold tolerance (T_crit-cold_)*.

For the PSII and heat tolerance assays, an array of 30 leaves was taped to paper and placed on the Peltier plate surface with heavy glass on top to maximise thermal transfer between the plate and leaves. Every second leaf had a calibrated type-T thermocouple attached to the underside of the leaf in a checkerboard pattern, which were interfaced with a DataTaker datalogger (DT80; Lontek, NSW, Australia) recording every 5 s, so that leaf temperature could be measured or approximated directly. The Peltier plate was held at 25°C and leaves were initially light-adapted under 280 µmol photons m^-2^ s^-1^ for 30 min to obtain stable minimal fluorescence (*F*_0_′) values, after which we applied three saturating pulses at ∼4,000 μmol photons m^-2^ s^-1^ for 720 ms each at 2 min intervals to determine the maximal fluorescence (*F*_M_′) when PSII reaction centres were closed. Variable fluorescence in the light (*F*_V_′ = *F*_M_′ – *F*_0_′) was then used to calculate φPSII (equivalent to *F*_V_′/*F*_M_′) (Baker, 2008) from the mean of the three measurements. These leaves were then dark-adapted for 25 min to reach a stable minimal chlorophyll fluorescence in the dark (*F*_0_) prior to applying a single saturating pulse to measure *F*_V_/*F*_M_, which is the equivalent of φPSII in the dark (Baker, 2008). After 1 min, we began recording *F*_0_ values continuously at 20 s intervals using a weak blue pulse modulated measuring light (0.5 μmol photons m^-2^ s^-1^) at 1 Hz to measure chlorophyll fluorescence without driving PSII photochemistry. We measured the *T-F*_0_ while heating the Peltier plate to 65°C at a rate of 30°C h^-1^.

Another similar system was used to measure cold tolerance limits in a cool room set to 6±2°C. The system was the same as described above except all 30 leaves had thermocouples attached to leaf undersides and the Peltier plate was set to 6°C to match the ambient cool room temperature. Leaves were dark adapted for 25 min before the temperature was reduced to –20°C at 15°C h^-1^, while recording *F*_0_ values continuously at 20 s intervals, and thermocouple (leaf) temperature every 5 s with a DataTaker datalogger (DT85-S4).

The *T-F*_0_ curve shows a stable, slow rise of *F*_0_ values until a critical temperature threshold occurs where *F*_0_ rises rapidly. The critical limit for both heat tolerance (*T*_crit-hot_) and cold tolerance (*T*_crit-cold_) is the temperature at the inflection point of intersecting regression lines for each of the *T-F*_0_ rise phases of relative *F*_0_, which is used as an indication of the thermal tolerance limits of photochemistry (Arnold *et al*., 2021). In both cases, the inflection point was calculated using a break-point regression analysis of the leaf temperature and relative *F*_0_ values using the *segmented* R package (Muggeo, 2017) in the R Environment for Statistical Computing v4.3.1 (R Core Team, 2023).

### The hailstorm and measurements at harvest

Unfortunately, on 20 January 2020, one of the most severe hailstorms in recorded history in Canberra damaged the glasshouses containing the plants (Rickards and Watson, 2020). There was no direct damage to the study plants because a shade screen was present, but the two controlled temperature treatments were lost due to damaged glasshouse infrastructure.

Cooling systems were restored 21 January 2020, but heating could not be restored. To ensure we did not therefore lose the investment in the long-term experiment, we took measurements on all plants immediately following the hailstorm. Specifically, we measured *chlorophyll content* using a handheld chlorophyll meter (SPAD-502; Konica Minolta Inc., Osaka, Japan) and *leaf mass per area (LMA)* on all plants within four days following the storm. Three healthy leaves were removed from each plant, immediately measured for chlorophyll content, and then scanned for leaf area before being placed in a drying oven at 60°C for > 72 h for weighing and calculating LMA. We had initially planned to continue the experiment until autumn (a further eight weeks), reduce temperatures to induce senescence, and then measure lifetime fitness and biomass. However, since controlled senescence was not possible, and since different plants were at various stages of flowering and producing seed at the time of the storm, we used a proxy for lifetime fitness across all plants: the *total number of reproductive stems* on an individual (i.e., flowers, capsules, and hardened, brown stems that clearly indicated a capsule had matured on the stem). We confirmed that the total number of reproductive stems was a suitable index of fitness by estimating its correlation with the harvested capsule mass × number of capsules weighed from a representative subset of 100 plants. We found a strong correlation (Pearson’s *r* = 0.873 ± 0.056), which did not differ among treatments (Supplementary Table S2, Supplementary Fig. S2). On 24-25 January 2020, we counted the number of reproductive stems on all plants, as well as collecting mature capsules. However, due to the hailstorm alternative controlled temperature growth space was limited and we elected to systematically harvested half of all plants (blocks 2 and 4) following the reproductive stem count. We harvested these plants and measured *dry above-ground biomass* in bags for drying at 60°C for > 72 h and subsequent weighing.

Plants from the blocks that were not harvested immediately following the hailstorm (blocks 1 and 3) were moved to four controlled environment Growth Capsules (Photon Systems Instruments, Brno, Czech Republic) run by the Australian Plant Phenomics Network at ANU where we aimed to continue the experiment. The Growth Capsules were set to match the glasshouse conditions as best possible, however the plants did not thrive in the Growth Capsules due to lower light, reduced air flow, and higher humidity. After 10 days in Growth Capsule conditions, we therefore measured thermal tolerance and chlorophyll fluorescence traits on the plants that had not yet been measured, and then ceased the experiment on 13-14 February 2020. We repeated the reproductive stem count and added new stems to the count from 24-25 January 2020, and then harvested these plants for biomass as above.

The sample sizes for each trait were *n* = 1,023 for number of reproductive stems, *n* = 1,003 for flowering onset, *n* = 988 for above-ground biomass, *n* = 984 for chlorophyll content, *n* = 975 for LMA, *n* = 717 for φPSII, *n* = 717 for *F*_V_/*F*_M_, *n* = 685 for *T*_crit-hot_, and *n* = 707 for *T*_crit-cold_. The inherent differences between the pre- and post-hailstorm measurements and harvesting are explicitly accounted for in our statistical analyses.

### Statistical analyses

For all analyses of the F3 phenotypic traits, models were fit using the R package *brms* (Bürkner, 2018) in the R Environment for Statistical Computing v4.3.1 (R Core Team, 2023). All *brm* models were run using four chains, each with 4000 iterations, 2000 of which were sampling, with *adapt_delta* ≥ 0.99 and *max_treedepth* = 15 so that the majority of R≤ 1.005, indicating that chains had effectively mixed. All response variable distributions exhibited some skewness, therefore we set *skew_normal* distributions for the univariate *brm* models, which are an extension of the normal (Gaussian) distribution family that also estimate a skew parameter. We verified that skew-normal models were a good fit to the data and that they were a better fit than models using a Gaussian distribution with posterior predictive checking (Gabry *et al*., 2019). To facilitate model convergence, φPSII and *F*_V_/*F*_M_ were both scaled by a factor of ten to avoid very small parameter estimates.

To test the main effects of growth temperature, parental temperature, and inbreeding on each trait and its plasticity, we initially fit univariate random regression mixed models (RRMMs; Arnold *et al*., 2019a) that included a structured pedigree (often called an ‘animal model’; Kruuk, 2004; Wilson *et al*., 2010), following the R *brms* form:

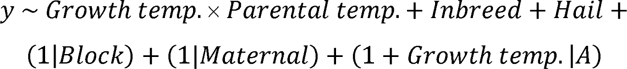

*y* is the phenotypic trait. Fixed effects were *Growth temp.*, a two-level factor of growth temperature; *Parental temp.*, a two-level factor of parental temperature; and their interaction; *Inbreed*, a two-level factor of inbreeding (outcrossed or self-pollinated); and *Hail*, a two-level factor of whether the measurement was taken before or after the hailstorm. Random intercepts were: *Block*, a four-level factor of experimental block; *Maternal*, an identification term for the F2 ovule donor to quantify maternal effects with 102 levels; and *A*, the additive genetic component with covariance structure defined by a pedigree of relatedness values among individuals, which was converted into an inverse *A* matrix using the *MCMCglmm* package (Hadfield, 2010). We included the random slope term *Growth temp.* with the *A* term to test for a genotype × environment (G×E) interaction. To evaluate whether the G×E term was important, we compared models with and without the *Growth temp.* slope term using leave-one-out cross validation (LOO-CV) to estimate predictive accuracy of each candidate model (Vehtari *et al*., 2017). We then calculated Bayesian stacking weights, which evaluate the average performance of the combined posterior predictive distribution of candidate models (Yao *et al*., 2018). We report the full model including the slope term given that our models were not overparameterized relative to our sample sizes. We include *R*^2^ values for mixed-effects models: marginal *R*^2^ (m*R*^2^) to estimate variance explained by fixed effects and the difference between m*R*^2^ and conditional *R*^2^ (c*R*^2^) to estimate variance explained by random effects (Nakagawa and Schielzeth, 2013) using the *performance* package (Lüdecke *et al*., 2021).

To calculate narrow-sense heritability *h*^2^, we took the posterior distribution of the additive genetic variance *V*_A_ from the animal model and divided it by the total phenotypic variance *V*_P_, where *V*_P_ = (*V*_A_ + *V*_B_ *+ V*_M_ + *V*_R_), and *V*_B_ is block variance, *V*_M_ is maternal variance, and *V*_R_ is residual variance. Since we included a maternal effect term in the models, we also estimated the contribution of direct maternal effects *m*^2^ as *V*_M_/*V*_P_.

To test for linear (directional) and quadratic (stabilising or disruptive) selection on traits, we fit multiple regression models of the trait and fitness, similarly to Noble *et al*. (2013). We estimated standardised selection gradients by converting the number of reproductive stems, which was our proxy for fitness, to relative fitness (*w*, by dividing by the mean of each growth treatment) and each trait was mean-centred and standardised to unit variance (Lande and Arnold, 1983), for the overall (all plants), cool-grown plants, and warm-grown plants separately. For the overall model we also included linear and non-linear interaction terms with the trait and temperature to determine if selection varied with temperature. Linear selection gradients (β) came from regression models without quadratic and the growth temperature × parental temperature interaction terms, whereas quadratic selection gradients (γ) come from models including these terms (Lande and Arnold, 1983).

Quadratic terms and their 95% credible intervals (95% CI) were doubled prior to reporting, such that they can be interpreted as stabilising or disruptive selection gradients (Stinchcombe *et al*., 2008). Positive linear selection coefficients can be interpreted as directional selection where individuals with larger phenotypic trait values have higher relative fitness on average. Positive quadratic selection coefficients can be interpreted as individuals with trait values at the edges of the trait distribution have higher relative fitness on average (convex function shape; disruptive selection). Negative quadratic selection coefficients can be interpreted as individuals with trait values in the centre of the trait distribution have higher relative fitness on average (concave shape; stabilising selection). A trait that has both significant linear and quadratic coefficients indicates an overarching directional change with a non-linear shape.

### Note on the effects of the hailstorm and later harvest date or later trait measurements

The impact of the hailstorm (which caused a delay in measurement of some traits and later harvest time for a subset of plants) was evident in some traits. Importantly, all models in our analyses included a term to account for this effect. As expected, plants that were harvested later had a greater number of reproductive stems and greater biomass than plants harvested immediately after the hailstorm. The plants that had traits measured post-hailstorm also had slightly higher chlorophyll content and *F*_V_/*F*_M_, but φPSII and the thermal tolerance traits (*T*_crit-hot_ and *T*_crit-cold_) were unaffected by the hailstorm (Tables 1 and 2).

**Table 1:** Model output summaries for five phenotypic traits representing reproductive fitness, phenology, biomass, and two leaf traits.

| *Response variable:* | Total reproductive stems<br>(*n* = 989) | Flowering onset<br>(*n* = 1,003) | Dry biomass<br>(*n* = 987) | Chlorophyll content<br>(*n* = 983) | LMA<br>(*n* = 974) |
| --- | --- | --- | --- | --- | --- |
| *Fixed effects* |  |  |  |  |  |
| *Estimate [95% CI]* |  |  |  |  |  |
| Intercept (cool Growth temp., cool Parental temp., outcrossed) | **35.907 [28.982, 42.255]** | **45.421 [43.721, 47.094]** | **7.550 [6.732, 8.343]** | **32.910 [31.322, 34.666]** | **1.044 [0.946, 1.134]** |
| Growth temp. (warm) | **3.946 [0.843, 7.060]** | **-7.899 [-8.861, -6.913]** | **0.745 [0.173, 1.344]** | **3.186 [1.908, 4.436]** | **0.100 [0.047, 0.155]** |
| Parental temp. (warm) | -1.393 [-5.438, 2.449] | 1.153 [-0.336, 2.676] | 0.104 [-0.561, 0.758] | -0.204 [-1.468, 1.074] | 0.025 [-0.024, 0.073] |
| Growth temp. × Parental temp. | -0.543 [-4.781, 3.743] | -0.592 [-1.866, 0.626] | -0.153 [-0.915, 0.608] | 0.265 [-1.282, 1.816] | -0.059 [-0.127, 0.011] |
| Cross type (self-pollinated) | **-7.830 [-10.957, -4.789]** | **1.492 [0.458, 2.525]** | **-1.297 [-1.833, -0.744]** | 0.785 [-0.227, 1.793] | -0.002 [-0.041, 0.037] |
| Harvest date (later) | **6.496 [4.119, 8.877]** | -- | **4.416 [4.007, 4.820]** | **1.173 [0.376, 1.941]** | **0.155 [0.120, 0.189]** |
| *Random effects: variance components* |  |  |  |  |  |
| *Estimate (SD) [95% CI]* |  |  |  |  |  |
| $V_B$ intercept (block) | 4.587 [1.216, 14.030] | 0.409 [0.012, 1.709] | 0.375 [0.020, 1.401] | 0.990 [0.085, 3.328] | 0.071 [0.018, 0.242] |
| $V_A$ intercept (additive genetic) | 4.271 [0.743, 7.317] | 2.808 [1.948, 3.588] | 0.787 [0.336, 1.205] | 1.416 [0.274, 2.297] | 0.028 [0.002, 0.063] |
| $V_A$ slope (G×E) * | 1.830 [0.070, 5.025] | 1.130 [0.232, 2.021] | 0.664 [0.179, 1.147] | 1.538 [0.219, 2.732] | 0.062 [0.014, 0.106] |
| $V_A$ intercept-slope correlation | -0.023 [-0.927, 0.929] | -0.793 [-0.993, -0.242] | 0.575 [-0.178, 0.984] | 0.124 [-0.678, 0.926] | 0.294 [-0.735, 0.967] |
| $V_M$ intercept (maternal) | 4.343 [0.744, 7.101] | 0.964 [0.044, 2.059] | 0.596 [0.051, 1.119] | 1.018 [0.072, 2.015] | 0.028 [0.002, 0.063] |
| $V_R$ (residual) | 21.415 [20.277, 22.567] | 5.268 [5.009, 5.546] | 2.788 [2.653, 2.930] | 5.663 [5.391, 5.949] | 0.285 [0.271, 0.301] |
| *Model stacking weights* |  |  |  |  |  |
| Model with G×E * | 0.000 | 0.396 | 0.453 | **0.594** | **0.716** |
| Model without G×E * | **1.000** | **0.604** | **0.547** | 0.406 | 0.284 |
| *R<sup>2</sup>* |  |  |  |  |  |
| mR<sup>2</sup> (fixed effects) | 0.247 | 0.582 | 0.546 | 0.228 | 0.026 |
| cR<sup>2</sup> - mR<sup>2</sup> (random effects) | 0.561 | 0.241 | 0.171 | 0.387 | 0.032 |

Estimates are posterior modes with [95% CIs]; bold represents fixed effects that have 95% CIs that are distinct from zero; Harvest date refers to plants that were harvested either immediately following the hailstorm or later, see *Methods* for details; \* Model outputs reported are full models that include G×E term, however please see the Model stacking weights for whether there is statistical support (bold) for the G×E term.

**Table 2:** Model output summaries for four phenotypic traits representing photosynthetic physiology and thermal tolerance.

| *Response variable:* | $F_V/F_M$<br>(*n* = 717) | $\phi\text{PSII}$<br>(*n* = 717) | $T_{\text{crit-hot}} (^{\circ}\text{C})$<br>(*n* = 685) | $T_{\text{crit-cold}} (^{\circ}\text{C})$<br>(*n* = 707) |
| --- | --- | --- | --- | --- |
| *Fixed effects* |  |  |  |  |
| *Estimate [95% CI]* |  |  |  |  |
| Intercept (cool Growth temp., cool Parental temp., outcrossed) | **7.277 [6.969, 7.585]** | **2.574 [2.134, 2.981]** | **44.459 [43.201, 45.691]** | **-13.256 [-14.312, -12.301]** |
| Growth temp. (warm) | **0.089 [0.029, 0.150]** | **0.332 [0.097, 0.560]** | 0.256 [-0.158, 0.675] | 0.345 [-0.052, 0.750] |
| Parental temp. (warm) | 0.054 [-0.005, 0.115] | 0.037 [-0.196, 0.270] | -0.039 [-0.451, 0.376] | 0.186 [-0.217, 0.591] |
| Growth temp. × Parental temp. | -0.033 [-0.116, 0.047] | -0.023 [-0.339, 0.290] | 0.059 [-0.503, 0.627] | -0.158 [-0.720, 0.398] |
| Cross type (self-pollinated) | 0.036 [-0.008, 0.082] | 0.082 [-0.091, 0.254] | 0.003 [-0.316, 0.326] | -0.186 [-0.490, 0.119] |
| Hail (measured post-hail) | **-0.124 [-0.221, -0.031]** | -0.028 [-0.312, 0.253] | 0.395 [-0.208, 1.034] | -0.101 [-0.623, 0.491] |
| *Random effects: variance components* |  |  |  |  |
| *Estimate (SD) [95% CI]* |  |  |  |  |
| $V_B$ intercept (block) | 0.182 [0.008, 0.938] | 0.245 [0.007, 1.167] | 0.880 [0.157, 3.091] | 0.663 [0.083, 2.414] |
| $V_A$ intercept (additive genetic) | 0.027 [0.001, 0.075] | 0.124 [0.006, 0.310] | 0.228 [0.012, 0.521] | 0.182 [0.007, 0.462] |
| $V_A$ slope (G×E) * | 0.042 [0.002, 0.110] | 0.174 [0.007, 0.423] | 0.307 [0.017, 0.696] | 0.334 [0.025, 0.710] |
| $V_A$ intercept-slope correlation | -0.268 [-0.976, 0.902] | -0.201 [-0.969, 0.902] | -0.136 [-0.951, 0.921] | -0.083 [-0.942, 0.932] |
| $V_M$ intercept (maternal) | 0.022 [0.001, 0.059] | 0.117 [0.006, 0.267] | 0.203 [0.009, 0.462] | 0.141 [0.007, 0.367] |
| $V_R$ (residual) | 0.336 [0.317, 0.358] | 1.015 [0.958, 1.074] | 1.793 [1.696, 1.901] | 2.252 [2.123, 2.390] |
| *Model stacking weights* |  |  |  |  |
| Model with G×E * | 0.041 | 0.001 | 0.000 | 0.309 |
| Model without G×E * | **0.959** | **0.999** | **1.000** | **0.691** |
| *R<sup>2</sup>* |  |  |  |  |
| mR<sup>2</sup> (fixed effects) | 0.011 | 0.023 | 0.018 | 0.011 |
| cR<sup>2</sup> - mR<sup>2</sup> (random effects) | 0.094 | 0.094 | 0.335 | 0.199 |

Estimates are posterior modes with [95% CIs]; bold represents fixed effects that have 95% CIs that are distinct from zero; Hail refers to plants that were measured for these traits before or after the hailstorm, see *Methods* for details; \* Model outputs reported are full models that include G × E term, however please see the Model stacking weights for whether there is statistical support (bold) for the G × E term.

## Results

### Q1: To what extent do a suite of phenotypic traits respond to growth temperature through phenotypic plasticity?

To test for plasticity in trait responses to temperature, we compared the change in mean trait value between cool and warm growth temperatures among parental temperature and cross type (self-pollinated *vs* outcrossed) groups. We present the overall mean effects in Fig. 1 and mean treatment- and family-level reaction norms in Supplementary Fig. S3, as well as summary statistics for each trait in Supplementary Table S1. Warm growth temperature had a significant positive effect on reproductive fitness, biomass, chlorophyll content, LMA, *F*_V_/*F*_M_, and φPSII (Fig. 1A,C-G, Tables 1, 2). Flowering onset also occurred significantly earlier (7.8 days on average) in warm-grown plants (Fig. 1B, Table 1). Seven traits showed significant phenotypic plasticity to growth temperature, with extensive variation around these average effects (Supplementary Fig. S3). There was no evidence of growth temperature effects on either heat or cold tolerance of PSII. Although the warm-grown plants exhibited a slightly higher *T*_crit-hot_ than the cool-grown plants, as would be expected with a thermal acclimation response, this difference was not significant and the average response was canalised (Fig. 1H, Table 2). Similarly, *T*_crit-cold_ did not differ significantly between treatments and was also, on average, canalised (Fig. 1I, Table 2).

**Fig. 1:**
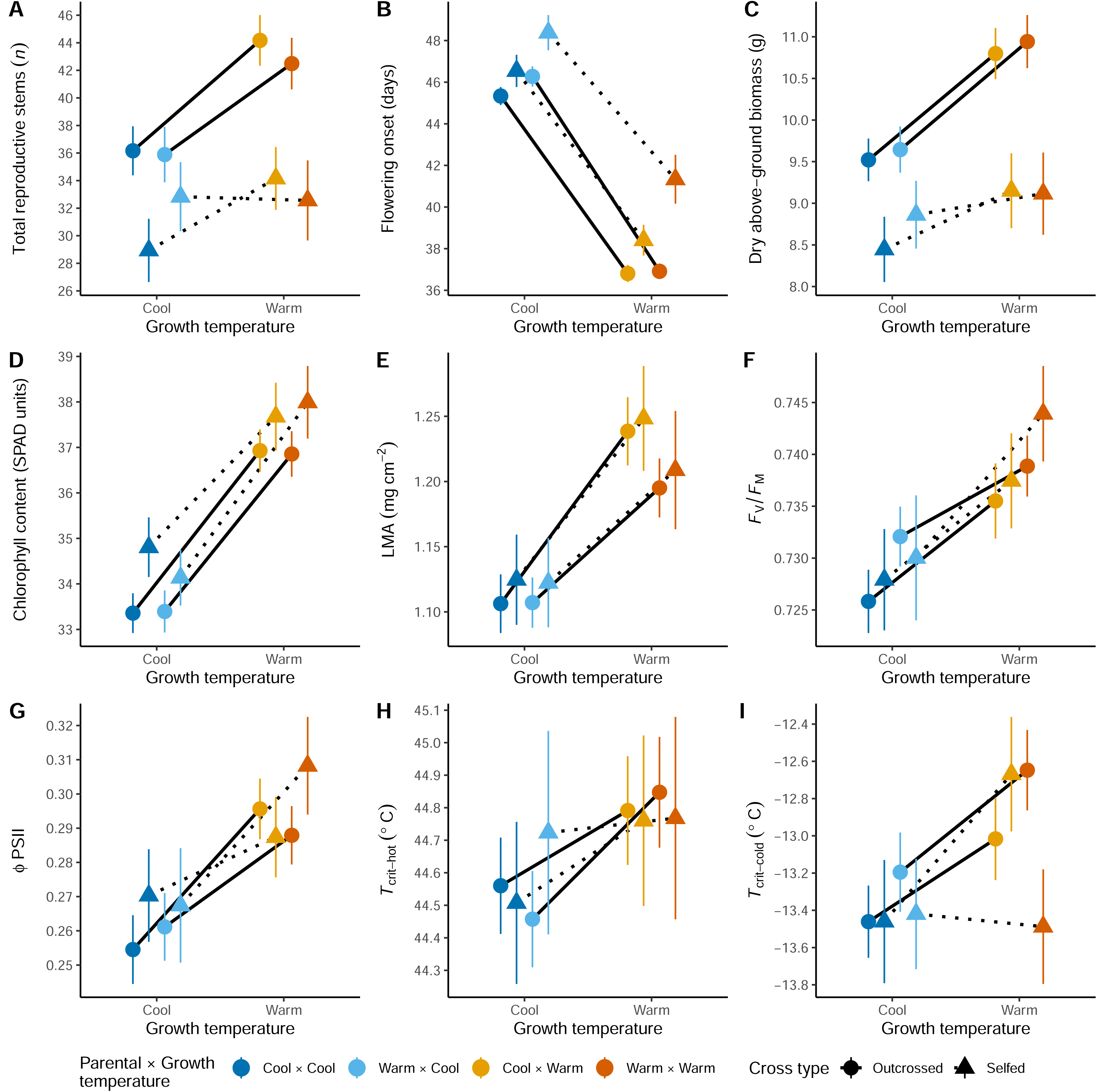
Mean population-level reaction norms of phenotypic traits: (A) total reproductive stems, (B) flowering onset, (C) biomass, (D) chlorophyll content, (E) LMA, as well as photosystem traits: (F) *F*_V_/*F*_M_ and (G) φPSII, and thermal tolerance traits: (H) *T*_crit-hot_ and (I) *T*_crit-cold_ in response to growth temperature treatments. Within each cool and warm growth temperature treatment, plants were grown under an environment that was either cool (blues) or warm (oranges) and were offspring plants were from either outcrossed (solid lines) or self-pollinated (dotted lines) parents. Each parental × growth temperature combination is coloured as follows: parental plants grown under a cool environment that had offspring grown in *i*) a cool environment (dark blue) or *ii*) a warm environment (light orange), and parental plants grown under a warm environment that had offspring grown in *iii*) a cool environment (light blue) or *iv*) a warm environment (dark orange). Reaction norms are drawn based on connections between a shared parental environment and cross type (e.g., parental cool × growth cool and outcrossed is connected to parental cool × growth warm and outcrossed). Points and error bars represent means ± S.E. of the raw data.

### Q2: Is there any evidence of transgenerational plasticity or benefits for offspring that are grown under conditions that match the conditions in which their parents were grown?

In this experiment, we used a factorial design to separate the effects of growing parental plants under relatively cool and warm temperatures and the subsequent effects of their offspring growing under the same (matched) or opposite (mismatched) temperature regimes. We hypothesised that development of seed under the parental temperature that matched the offspring growth temperature (e.g., cool × cool, or warm × warm) would exhibit phenotypes that performed better than mismatched offspring (e.g., cool × warm, or warm × cool).

However, we found no evidence that the parental temperature had any effect on any of the measured traits (Fig. 1, Supplementary Fig. S4), nor of any significant interactions between growth and parental temperatures (Tables 1 and 2). Therefore, there was no evidence in any trait that matching parent-offspring environments was beneficial, nor was there any evidence that mismatching was detrimental (Supplementary Figs S5 and S6).

### Q3: Are the phenotypic traits heritable, and are there either maternal effects or G×E interactions?

Estimates of heritability were relatively low across all traits (ranging from 0.01 to 0.14), with flowering onset, biomass, and chlorophyll content being the only traits for which there was support for a non-zero heritability (Table 3). While maternal effects were included in all models, estimates of their variance components were small and the credible intervals of the estimates were not clearly distinct from zero (Table 3).

**Table 3:** Summary of heritability and maternal effects on each phenotypic trait.

| *Phenotypic trait* | $h^2$ ($V_A/V_P$) | $m^2$ ($V_M/V_P$) |
| --- | --- | --- |
| Total reproductive stems | 0.040 [$<0.001$, 0.085] | 0.040 [$<0.001$, 0.085] |
| Flowering onset | **0.142 [0.056, 0.226]** | 0.050 [$<0.001$, 0.136] |
| Biomass | **0.114 [0.043, 0.189]** | 0.042 [$<0.001$, 0.107] |
| Chlorophyll content | **0.078 [0.018, 0.137]** | 0.029 [$<0.001$, 0.082] |
| LMA | 0.025 [$<0.001$, 0.054] | 0.015 [$<0.001$, 0.045] |
| $F_V/F_M$ | 0.005 [$<0.001$, 0.019] | 0.005 [$<0.001$, 0.020] |
| $\phi\text{PSII}$ | 0.016 [$<0.001$, 0.052] | 0.016 [$<0.001$, 0.051] |
| $T_{\text{crit-hot}}$ | 0.017 [$<0.001$, 0.051] | 0.014 [$<0.001$, 0.046] |
| $T_{\text{crit-cold}}$ | 0.010 [$<0.001$, 0.032] | 0.005 [$<0.001$, 0.020] |

Estimates are posterior modes with [95% CIs]. $V_A$ is additive genetic variance; $V_P$ is total phenotypic variance; $V_M$ is maternal variance. Note that *brms* models are parameterised such that variance components cannot be exactly zero. The minimum value for determining whether a credible interval is distinguishable from zero is 0.001. As such, any value $> 0.001$ (shown in bold) indicates 95% CIs that are clearly distinct from zero, providing evidence for either heritability or maternal effects on that phenotypic trait.

Flowering onset had a clear negative correlation between *V*_A_ intercept and slope, indicating that families that flowered early on average were those with the lowest plasticity. However, we only found support for G×E (here specifically referring to *V*_A_ depending on growth environment) being important for two traits: chlorophyll content and LMA (Table 1).

### Q4: What is the direction and strength of selection on the traits and how does it vary with growth temperature?

Our estimates of selection varied substantially among the traits. We found evidence for both linear (β; directional) and non-linear (γ; stabilising or disruptive) selection gradients for flowering onset, biomass, and LMA (Table 4; Fig. 2), but no clear evidence of selection on any other phenotypic trait (Table 4).

**Fig. 2:**
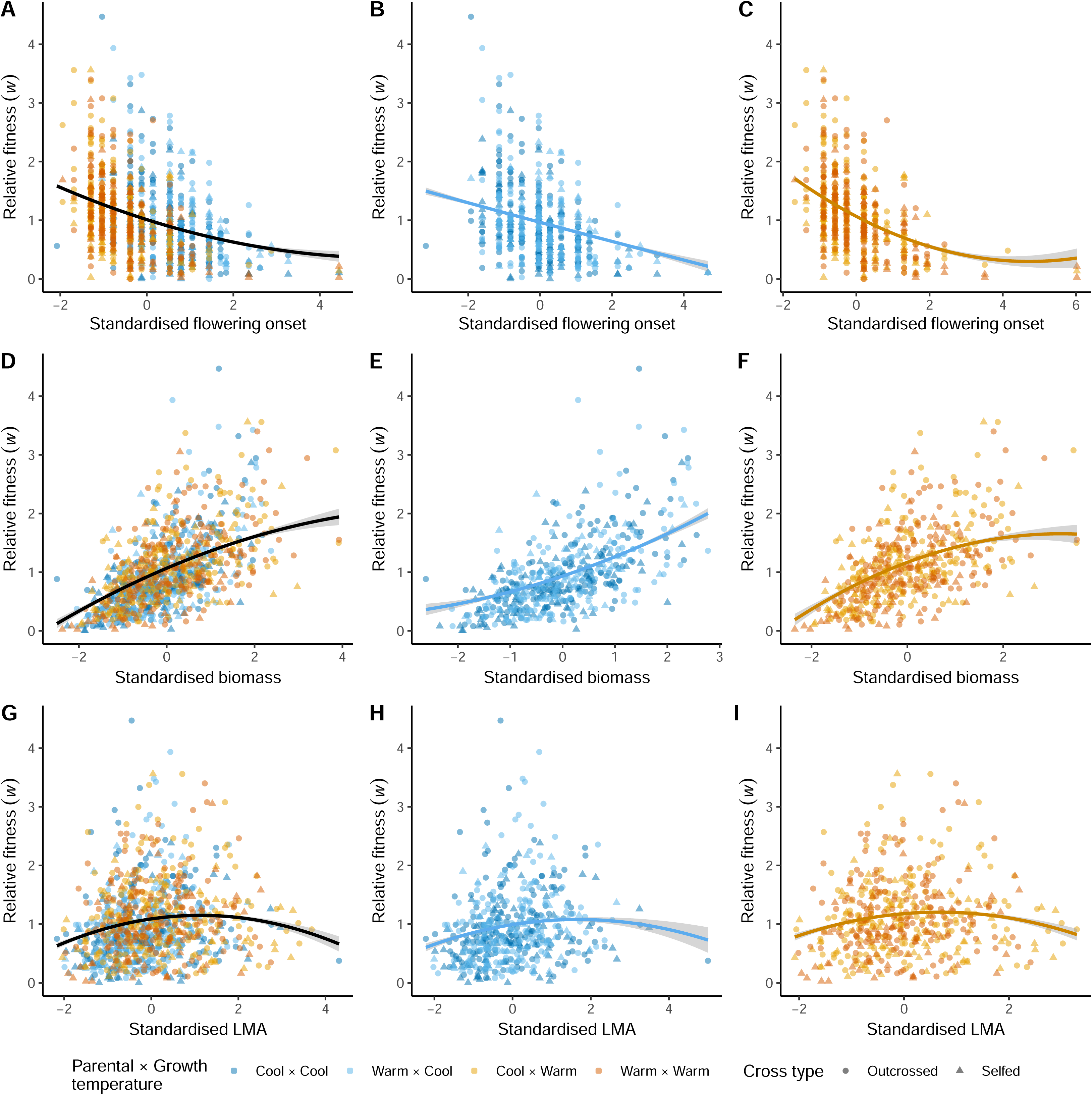
Relationships between relative fitness (*w*; calculated as the number of reproductive stems produced divided by the mean of each growth treatment) and three standardised (within each temperature treatment) phenotypic traits (A-C: flowering onset, D-F: biomass, G-I: LMA) that demonstrated non-zero selection. Left panels are all plants combined, middle panels are cool-grown plants, and right panels are warm-grown plants. Offspring plants were either outcrossed (circles) or self-pollinated (triangles) and were derived from parents that had a parental environment that was either warm or cool. The overall quadratic model fit (posterior predictions) is plotted on each panel. Note that scaling for standardising trait values on the *x*-axis is applied in each data subset and therefore individual data point positions differ along the *x*-axis between all plants, cool-grown plants, and warm-grown plants. Linear (β) and quadratic (γ) selection coefficients and 95% CIs are given in Table 4.

**Table 4:**
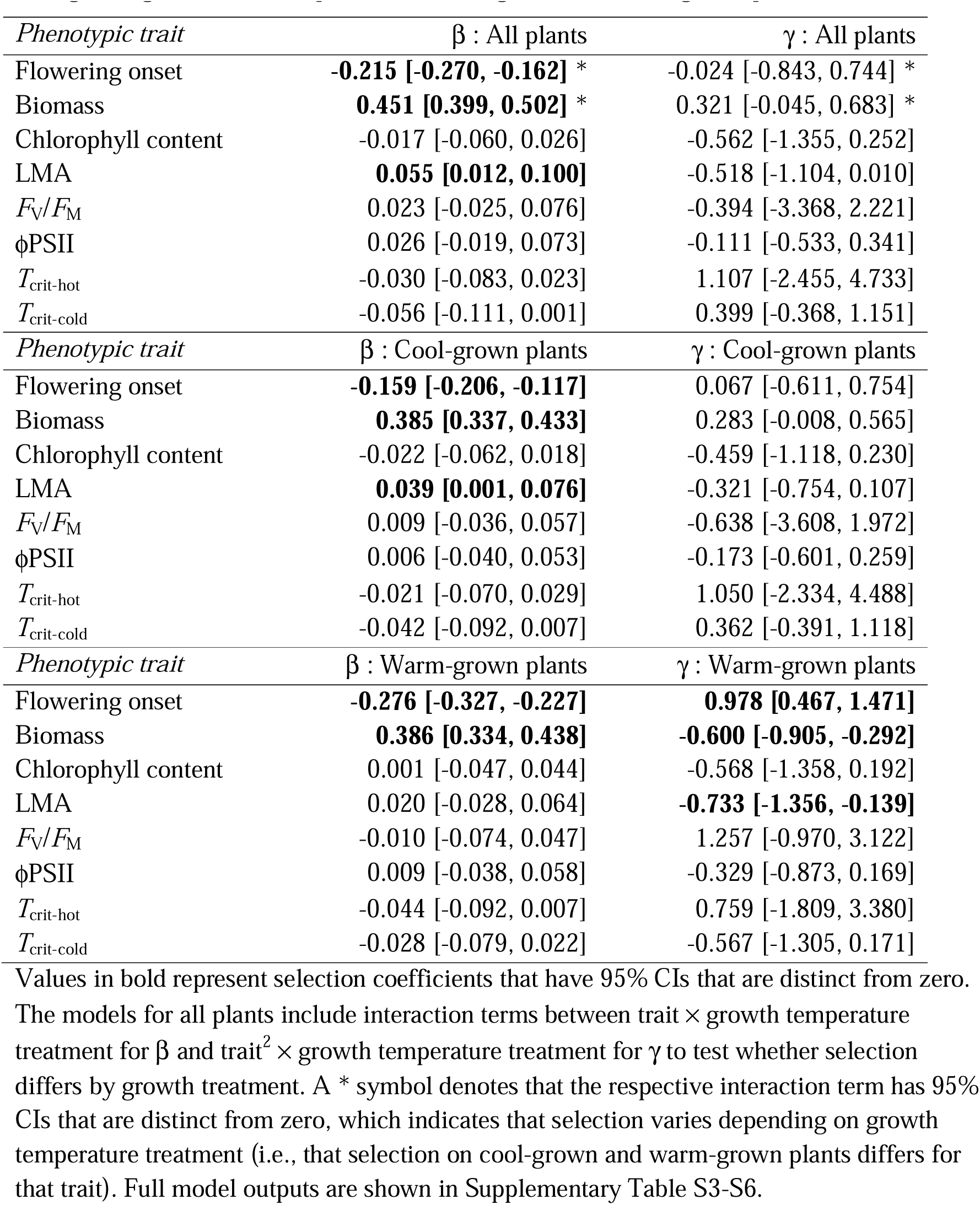
Linear (β) and quadratic (γ) selection coefficients on each of the phenotypic traits (excluding total reproductive stems because it is used to calculated relative fitness) under both growing conditions (all plants), and cool-grown and warm-grown plants alone.

| *Phenotypic trait* | $\beta$ : All plants | $\gamma$ : All plants |
| --- | --- | --- |
| Flowering onset | **-0.215 [-0.270, -0.162] \*** | -0.024 [-0.843, 0.744] * |
| Biomass | **0.451 [0.399, 0.502] \*** | 0.321 [-0.045, 0.683] * |
| Chlorophyll content | -0.017 [-0.060, 0.026] | -0.562 [-1.355, 0.252] |
| LMA | **0.055 [0.012, 0.100]** | -0.518 [-1.104, 0.010] |
| $F_v/F_m$ | 0.023 [-0.025, 0.076] | -0.394 [-3.368, 2.221] |
| ϕPSII | 0.026 [-0.019, 0.073] | -0.111 [-0.533, 0.341] |
| $T_{crit-hot}$ | -0.030 [-0.083, 0.023] | 1.107 [-2.455, 4.733] |
| $T_{crit-cold}$ | -0.056 [-0.111, 0.001] | 0.399 [-0.368, 1.151] |
| *Phenotypic trait* | $\beta$ : Cool-grown plants | $\gamma$ : Cool-grown plants |
| Flowering onset | **-0.159 [-0.206, -0.117]** | 0.067 [-0.611, 0.754] |
| Biomass | **0.385 [0.337, 0.433]** | 0.283 [-0.008, 0.565] |
| Chlorophyll content | -0.022 [-0.062, 0.018] | -0.459 [-1.118, 0.230] |
| LMA | **0.039 [0.001, 0.076]** | -0.321 [-0.754, 0.107] |
| $F_v/F_m$ | 0.009 [-0.036, 0.057] | -0.638 [-3.608, 1.972] |
| ϕPSII | 0.006 [-0.040, 0.053] | -0.173 [-0.601, 0.259] |
| $T_{crit-hot}$ | -0.021 [-0.070, 0.029] | 1.050 [-2.334, 4.488] |
| $T_{crit-cold}$ | -0.042 [-0.092, 0.007] | 0.362 [-0.391, 1.118] |
| *Phenotypic trait* | $\beta$ : Warm-grown plants | $\gamma$ : Warm-grown plants |
| Flowering onset | **-0.276 [-0.327, -0.227]** | **0.978 [0.467, 1.471]** |
| Biomass | **0.386 [0.334, 0.438]** | **-0.600 [-0.905, -0.292]** |
| Chlorophyll content | 0.001 [-0.047, 0.044] | -0.568 [-1.358, 0.192] |
| LMA | 0.020 [-0.028, 0.064] | **-0.733 [-1.356, -0.139]** |
| $F_v/F_m$ | -0.010 [-0.074, 0.047] | 1.257 [-0.970, 3.122] |
| ϕPSII | 0.009 [-0.038, 0.058] | -0.329 [-0.873, 0.169] |
| $T_{crit-hot}$ | -0.044 [-0.092, 0.007] | 0.759 [-1.809, 3.380] |
| $T_{crit-cold}$ | -0.028 [-0.079, 0.022] | -0.567 [-1.305, 0.171] |

Values in bold represent selection coefficients that have 95% CIs that are distinct from zero.

Relative fitness was higher in individuals with earlier flowering onset: individuals flowering later had generally very low fitness (Fig. 2A). The flowering onset of warm-grown plants had a stronger signal of selection than cool-grown plants (i.e., a more negative linear selection coefficient and a large quadratic coefficient; Table 4; Fig. 2B,C), where relative fitness was lower in warm-grown plants that had intermediate to high values of flowering onset (Fig. 2C; see Supplementary Tables S3–S6 for full models including the interaction between selection and temperature). Relative fitness was lowest in low biomass individuals, but there was a strong positive linear selection coefficient (increased fitness as biomass increased) for all plants combined and under both growing temperatures separately (Table 4; Fig. 2D,E).

Selection patterns differed between cool-grown and warm-grown plants (Supplementary Tables S3 and S5). Although there were no significant non-linear selection patterns in cool-grown plants (Table 4), we present the predicted non-linear fits for direct comparison to the warm-grown plants (Fig. 2). In warm-grown plants, there was a significantly negative quadratic selection coefficient for biomass, where the relationship between relative fitness and biomass tapered off at very high values of biomass (Fig. 2F). LMA was under stabilising selection for all plants combined, such that plants with intermediate values of LMA had higher relative fitness (Table 4; Fig. 2G-I). Cool-grown plants had a relatively small positive linear selection coefficient and a non-significant negative quadratic coefficient (Fig. 2H), whereas warm-grown plants were not under linear selection but showed a stabilising selection response that favoured intermediate LMA values (Fig. 2I).

### Q5: Are there effects of inbreeding on the suite of traits, and does any inbreeding depression vary with temperature?

We predicted that plants that were the result of outcrossing as opposed to self-pollination would have higher fitness due to inbreeding depression in the latter. Self-pollination had a significant negative effect on reproductive fitness (Table 1), such that these plants produced a mean of 7.7 fewer reproductive stems than plants that were outcrossed (19.3% reduction; Fig. 3A). Self-pollination delayed mean flowering onset by 2.1 days (5.1% reduction; Fig. 3B) and reduced mean biomass by 1.35 g (13.2% reduction; Fig. 3C). None of the other leaf traits, photosystem or thermal tolerance traits were affected significantly by self-pollination, and the effects of inbreeding depression were consistent across both growth temperatures (Supplementary Figs S7 and S8, Tables 1 and 2).

**Fig. 3.**
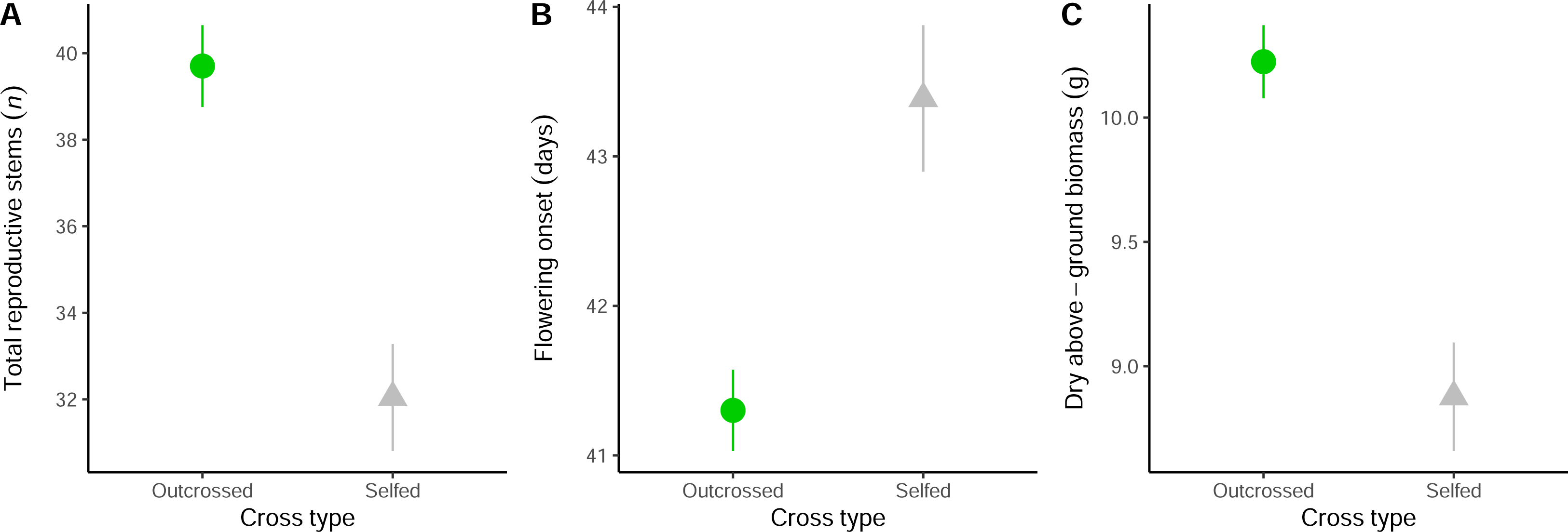
Tests for inbreeding depression. Mean differences between outcrossed and self-pollinated plants for fitness and phenotypic traits: (A) total reproductive stems, (B) flowering onset, and (C) biomass. Points and error bars represent means ± S.E. of the raw data. All phenotypic traits are shown in Supplementary Fig. S7. Model output is shown in Table 1.

## Discussion

In this study we tested the effects of growth temperature, parental temperature, and inbreeding on the multivariate phenotypes of an alpine plant with a mixed-mating system. We found strong phenotypic plasticity for most traits, with even 10°C warmer average growth temperatures having largely positive effects on fitness. There was substantial among-family variation in trait values in each environment, as well as in the direction and magnitude of reaction norms. Coupled with strong selection gradients and heritability of flowering onset, biomass, and chlorophyll content, we have evidence for plasticity in response to climate warming as well as evolutionary responses in *W. ceracea*, with limited indication that plasticity itself is adaptive.

### Growth temperature induces plastic responses in all traits except thermal tolerance

Phenotypic plasticity is a nearly ubiquitous response to warming conditions for functional traits that are limited by thermally-dependent reaction rates, or for traits that respond to abiotic cues associated with seasonal changes, such as photoperiod and temperature (Stotz *et al*., 2021). While prolonged or chronic warming certainly can be limiting for plants (Nievola *et al*., 2017; Lippmann *et al*., 2019), alpine plant growth and reproduction is typically restricted to a relatively short growing season that follows the release from cold temperature constraints (Körner, 2003; Dolezal *et al*., 2020). It is therefore reasonable to propose that in the non-limiting, well-watered, and warmer growing conditions, *W. ceracea* was stimulated to both grow and reproduce more than the cool-grown plants. This supports our hypothesised responses, except for that of heat tolerance. Warm-grown plants that had higher chlorophyll content and LMA, coupled with higher photosynthetic efficiency traits clearly allowed higher biomass production than cool-grown plants. Then, faster development and growth under warm conditions permitted earlier flowering onset that also increased the length of the reproductive period while allowing greater investment in reproduction. Our results indicate that these plastic responses are inducing an adaptive shift in the direction of higher fitness (Radchuk *et al*., 2019), and our results are consistent with empirical field research that finds warming in colder climate plant species stimulates growth and reproduction (Dolezal *et al*., 2020). For example, temperature enhancement using open top chambers in the field in Germany stimulated both growth and reproduction significantly during the growing season in alpine grassland species, although herbaceous perennials were less responsive than graminoids or shrubs (Kudernatsch *et al*., 2008).

The capacity for plants to increase their physiological thermal tolerance has been predicted to be a key response to climate warming (Geange *et al*., 2021). For example, *T*_crit-hot_ is well known to increase rapidly by 4°C or more within hours to days during an acute heat stress event (e.g., Andrew *et al*., 2023; Zhu *et al*., 2024). Long-term exposure to warm growth temperatures can also increase *T*_crit-hot_, for example by ∼0.16°C per 1°C of growth temperature (Zhu *et al*., 2018). We have previously observed plasticity in heat tolerance (*T*_crit-hot_) in the F1 and F2 generations of our *W. ceracea* experiments to long-term warming of 28-30°C. In those experiments, warm-grown plants increased their critical heat tolerance limits by 0.7-3.2°C relative to cool growing conditions (Notarnicola *et al*., 2021; Arnold *et al*., 2022), while also having a range of important effects on phenotypic and reproductive traits. It is worth noting that both these previous studies constrained pot sizes and used different, more confined controlled growth environments, which limits the value of making comparisons to our current glasshouse study (see Karitter *et al*., 2023 for a discussion of differences in phenotypic expression among common-environment experiments). We have also shown that moderate warming to 30°C can result in upregulation of genes related to post-transcriptional processes and downregulation of genes related to photosynthesis-related processes in *W. ceracea* (Notarnicola *et al*., 2023a). These suites of genetic changes that differ between cool to warm growing conditions may not correspond directly to changes in the phenotypic trait *T*_crit-hot_, which increased by a mean of only ∼0.26°C in the current study. Therefore, this canalisation of *T*_crit-hot_ to warming that we observed suggests that the warm-grown plants (while being well-watered) were not severely stressed, despite 30°C being far warmer than typical growing conditions for alpine *W. ceracea*. The warm treatment may have also alleviated temperature limitation on growth processes that can occur in alpine environments, which aligns with other trait responses that we observed. However, we encourage future studies to also embark on detailed examinations of thermal tolerance responses using thermal performance curves to determine pivotal temperature thresholds, as well as plasticity and selection thereof (Wooliver *et al*., 2022).

### Strong non-linear selection on heritable traits, especially under warm conditions

Warming has the potential to drive rapid evolutionary change in plant phenotypes, provided that phenotypic traits have a relationship with fitness and are therefore under selection, and that variation in the traits is heritable (Anderson and Song, 2020; Scheiner *et al*., 2020). Here our hypothesis that flowering onset, biomass, LMA, and heat tolerance would be heritable and under selection was partially supported. We found that flowering onset, biomass, and chlorophyll content were clearly heritable in *W. ceracea*. Chlorophyll content is proportional to the concentration of photosynthetic pigments and nitrogen in a leaf (Ling *et al*., 2011); it is heritable in wheat (Rosyara *et al*., 2010; Said *et al*., 2022), and has a relatively strong relationship with flowering in other crop species (Senger *et al*., 2014). To the best of our knowledge, there has not been another study reporting the heritability of chlorophyll content in a wild species, but our findings here suggest that variation in photosynthetic pigment concentration at least has a genetic basis. The low heritability of total reproductive stems (i.e., our best measure of fitness) may reflect depletion of genetic variance for fitness as expected from evolutionary theory (Falconer and Mackay, 1996; Kruuk *et al*., 2000), as well as large other sources of variance.

Here we found that both flowering onset and biomass were heritable as well as under relatively strong selection. There was significant negative directional selection (β) on flowering onset in all cases, where earlier flowering is favoured, and relative fitness in warm-grown plants also declined non-linearly (non-zero γ) with later flowering. These observed patterns of selection on flowering phenology align exactly with the findings from the relative cool and warm ends of the spectrum from a natural geothermal heating experiment on selection on flowering phenology in the short-lived perennial herb *Cerastium fontanum* in Iceland (Valdés *et al*., 2019). In our study, biomass was under strong positive directional selection across all environments, where larger plants had higher relative fitness, although fitness tapered off for larger warm-grown plants. Biomass can be a reasonable proxy for fitness (Younginger *et al*., 2017), where selection can favour larger individual size to facilitate plant performance (Aspi *et al*., 2003). Nevertheless, the plateau in relative fitness at larger sizes in warm-grown plants may be because the largest individuals would have relatively higher water demands and stronger resource allocation trade-offs than their smaller conspecifics under warming. Glasshouse studies can exacerbate effects of water limitation, and indeed we observed anecdotally that at their peak size, larger individuals in the warm-grown treatment began to wilt toward the end of hot, sunny days during the experiment. Heritability of biomass in these F3 plants aligns with previous findings of high *V*_A_ in both early growth rate and biomass in F2 *W. ceracea* plants (Arnold *et al*., 2022).

LMA was under positive directional selection overall and in cool-grown plants, but under stabilising selection in warm-grown plants, with intermediate to high but not extreme LMA values being favoured. LMA is an estimate of the density of carbon and nutrients in a set area of leaf tissue (i.e., the cost of tissue production for light interception) and is part of a trait complex that determines photosynthetic capacity, and nitrogen and water use efficiency (Poorter *et al*., 2009; Funk *et al*., 2021). Investment in high LMA may improve resource gain but only without critical water deficit (Ivanova *et al*., 2018), and potentially at the cost of reinvesting the acquired resources into vegetative rather than reproductive tissues (Flores *et al*., 2014). Thus, within a species, relatively low and high LMA values represent inefficient resource acquisition-use strategies that trade-off with reproduction, hence extreme values of LMA are selected against (Flores *et al*., 2014). Despite previously finding high intraspecific variation in *T*_crit-hot_ (Arnold *et al*., 2022), here it was neither heritable nor under selection, perhaps because photosynthetic heat tolerance does not affect fitness directly.

Taken together, our results suggest that traits contributing to light interception, growth, biomass, and flowering phenology are the key traits for ecological and evolutionary responses in plants to temperature. The capacity for these plants to tolerate and respond to warmer-than-typical temperatures demonstrates tolerance to moderate warming through a combination of phenotypic plasticity and intrinsic resilience. Growth, size, and reproductive traits respond to temperature over longer timescales (weeks to months) and contribute directly to fitness, whereas physiological traits regulate essential functions on shorter timescales (hours to days), but do not contribute directly to fitness. We highlight the need for future studies to take a demographic approach to studying plant responses to environmental stressors, integrating across the life cycle. Early life stages that are critical for establishment, growth, and survival, and reproductive stages that may be sensitive to temperature extremes and which directly affect fitness are typically less often studied than seeds or young adult plants. Finally, considering that warm-grown plants have different (non-linear) patterns of selection to cool-grown plants, future studies should concentrate on investigating novel or edge conditions to determine tipping points or sensitivity for ecological and evolutionary responses.

### No evidence of transgenerational plasticity via parental temperature or maternal effects

By applying a reciprocally matched-mismatched parent-offspring environments design, combined with the breeding structure, we could test for transgenerational plasticity via the parent environment effects and maternal effects through a pedigree (Uller *et al*., 2013). Evidence for matching parent-offspring environments benefitting offspring is relatively weak overall in plants (Uller *et al*., 2013). Based on earlier results with *W. ceracea* (Wang *et al*., 2021; Notarnicola *et al*., 2023b), we hypothesised that there could be a small benefit for offspring performance and fitness when matching their parent environment. However, we found no convincing evidence for any form of transgenerational plasticity, beneficial or not.

Using the same breeding design for F2 and F3 families as in the current experiment, Wang *et al*. (2021) tested for transgenerational plasticity in early life traits in *W. ceracea*. Seeds from parent plants grown in warm conditions had delayed germination (extended dormancy) and reduced germination success irrespective of their germination temperature, but none of these effects persisted to affect seedling growth (Wang *et al*., 2021). In a comprehensive reciprocal transplant experiment with *Boechera stricta* across an elevation gradient, Wadgymar *et al*. (2018a) found transgenerational plasticity in the early life traits of seed viability, germination, and dormancy. Transgenerational plasticity interplayed with within-generation plasticity across elevations and the effects of both were complex and context-specific, but parental environmental effects largely did not persist to later life (Wadgymar *et al*., 2018a). However, in a reciprocal environment experiment on *Lupinus angustifolius* under well-watered and drought stress treatments, Matesanz *et al*. (2022) found significant transgenerational plasticity that affected functional traits and reproduction of the offspring. Parental effects altered individual seed mass, flowering onset, and growth rate of the offspring, but these effects were not always beneficial, and offspring environment effects far outweighed the parental effects for specific leaf area, *F*_V_/*F*_M_, and lifetime reproductive output (Matesanz *et al*., 2022). Seed provisioning through maternal resource allocation affects seed viability, which in turn affects the probability of germination success (Haig and Westoby, 1988). Unsuitable, limiting, or stressful parental environments can also have direct adverse effects on reproductive tissues. For example, exposure to high temperature can disrupt reproductive development and reduce pollen viability, leading to smaller and/or less viable seeds, which have altered germination responses (Herman and Sultan, 2011; Sehgal *et al*., 2018; Goel *et al*., 2023).These examples highlight that transgenerational plasticity can certainly affect seed traits, but that persistent effects into adult phenotypes of the offspring might be less common or weaker (Herman and Sultan, 2011; Wang *et al*., 2021; Notarnicola *et al*., 2023b), although there are notable exceptions (e.g., Whittle *et al*., 2009).

### Inbreeding impairs reproduction and biomass but not physiological function

We predicted that inbreeding would significantly reduce plant performance and fitness compared to crossing, due to inbreeding depression. *Wahlenbergia ceracea* is protandrous and facultatively autogamous with a mixed-mating system (Nicotra *et al*., 2015), where self-pollination of the same flower is delayed by several days following flower opening, which provides reproductive assurance in the absence of external pollination (Goodwillie *et al*., 2005). Many alpine species with a mixed-mating strategy depend on external pollination to achieve their maximum potential seed set (Scheffknecht *et al*., 2007). It is therefore unsurprising that we found that self-pollination in *W. ceracea* caused marginally delayed flowering and slightly reduced total reproductive stems and biomass. The magnitude of the inbreeding depression effect on these fitness-related traits is comparable to the expected range from a meta-analysis of inbreeding effects on plant fitness (Angeloni *et al*., 2011). Inbreeding effects can be exacerbated in stressful environments (Armbruster and Reed, 2005), however we did not observe this effect, nor did we find any inbreeding effect on functional traits. We suggest that inbreeding (particularly in mixed-mating species) might affect fitness directly rather than indirectly through traits that mediate resource acquisition. However, we do not yet know whether negative effects of inbreeding on function and fitness would be exacerbated under more challenging conditions (e.g., heat coupled with drought) or extreme events (e.g., heatwaves).

### Conclusions and future directions

The capacity for plants to alter their phenotype in response to climate warming is frequently thought to be adaptive. Here we show through comprehensive analyses that in this alpine species, warming may alleviate restrictions on growth and reproduction, thereby improving fitness under warming through plasticity. The exception to this conclusion was thermal tolerance, which is likely already at sufficient levels. Only flowering onset and biomass were both heritable and clearly under selection, which differed between the cool and warm growth environments, and only chlorophyll content and LMA had any evidence for G×E. The effect of growth environment far exceeded any influence of parental environment; we found little evidence for substantial maternal effects or transgenerational plasticity in adult traits. Further, the effect of inbreeding by self-pollination was relatively small, providing reproductive assurance at low cost. We can conclude that the mixed-mating alpine herb *W. ceracea* clearly has capacity to respond rapidly to climate warming via phenotypic plasticity as well as the potential for evolutionary change across generations.

As the climate of alpine ecosystems changes, the duration of the growing season will extend, generating both new opportunities and new challenges for its inhabitants. Our experiment found that substantially warmer daytime temperatures (30°C) can still facilitate growth and reproduction in an alpine herb when water is not limiting. However, climate change is also expected to progressively dry some alpine ecosystems. Climate projections for some areas with seasonal snowpack have forecast reduced winter snowfall, earlier snowmelt in spring, and potentially decreases in summer and autumn precipitation events (Gobiet *et al*., 2014; Harris *et al*., 2016). Reductions in water supply have a clearly detrimental effect on most alpine plants (Sumner and Venn, 2021), and interactions between warming and water limitation are undoubtedly relevant for future climate scenarios in alpine plant communities (De Boeck *et al*., 2016; Winkler *et al*., 2016). Therefore, an essential next step in building an understanding of the importance of eco-evolutionary responses to climate change will be to test the role of water limitation in altering plastic and evolutionary responses to temperature.

Heat stress events are predicted to become more frequent, intense, and longer duration (Trancoso *et al*., 2020), on top of a background of mean climate warming (Harris *et al*., 2018). Extreme events have the potential to change fitness drastically and could be stronger selective events than gradual environmental change, which will alter evolutionary dynamics of populations in future (Gutschick and BassiriRad, 2003). The role of extreme events in the eco-evolutionary dynamics of alpine plants remains largely unexplored, despite alpine ecosystems being among the most vulnerable to and already impacted by climate change (Verrall and Pickering, 2020). Using genomic approaches to study climate change responses in natural populations could reveal the genomic architecture of traits exhibiting plasticity and under selection, improving our understanding of the mechanisms behind stress responses and their evolutionary potential (Notarnicola *et al*., 2023a). Employing a multifaceted research effort to strengthen our understanding of the roles of plasticity and evolutionary responses to realistic climate scenarios and extreme events is necessary to evaluate the potential for alpine plants to respond to future conditions.

## Supplementary data

*Table S1:* Summary statistics for all phenotypic traits.

*Table S2:* Linear regression model output testing fitness index suitability.

*Table S3:* Model outputs for linear selection by growth treatment 1.

*Table S4:* Model outputs for linear selection by growth treatment 2.

*Table S5:* Model outputs for quadratic selection by growth treatment 1.

*Table S6:* Model outputs for quadratic selection by growth treatment 2.

*Fig. S1:* Summary of the multi-generational breeding design.

*Fig. S2:* Correlation between total reproductive stems and reproductive mass.

*Fig. S3:* Reaction norms of phenotypic traits.

*Fig. S4:* Isolated parental temperature effects for phenotypic traits.

*Fig. S5:* Matched/mismatched parent-offspring environment effects on traits 1.

*Fig. S6:* Matched/mismatched parent-offspring environment effects on traits 2.

*Fig. S7:* Tests for inbreeding depression in phenotypic traits 1.

*Fig. S8:* Tests for inbreeding depression in phenotypic traits 2.

## Acknowledgements

We respectfully acknowledge the traditional custodians of the land from which plant seeds were originally collected and the land on which this research was conducted: the Ngunnawal, Ngambri, Ngarigo, and Walgalu people. We sincerely thank The Australian National University Plant Services staff: Steve Dempsey, Christine Larsen, Darren Marsh, Gavin Pritchard, and Jenny Rath for their extraordinary contributions to maintaining the glasshouse plants, and with hailstorm recovery efforts. We thank Richard Poiré for providing a second Imaging-PAM system and for offering space in the Growth Capsules after the hailstorm. The phenotyping and harvesting of the plants, particularly in the aftermath of the hailstorm, would not have been possible without the efforts of a substantial team of volunteers and colleagues, for whom we are very grateful: Monica Arnold, Verónica Briceño, Zachary Brown, Stephanie Courtney Jones, Jemimah Hamilton, Ophélie Lasne, Patrick Meir, Tenzin Norzin, Helen Osmond, Abigail Ryan, Natasha Salisbury, Alice Schacher, and Kaitlyn Spooner. Further thanks to Alexandra Catling, Kelli Gowland, Pamudika Kiridena, and Melissa Wakem for their contributions to generating the F3 lines.

## Author contributions

PAA – conceptualisation, experimental design, performed the experiments, analysed the data, wrote the first draft.

SW – performed the experiments, editing.

RFN – performed the experiments, editing.

ABN – conceptualisation, experimental design, funding acquisition, supervision, editing.

LEBK – conceptualisation, experimental design, funding acquisition, supervision, editing.

## Data availability

Data and R code will be available in the Dryad repository doi:10.5061/dryad.34tmpg4s9.

