## Supplementary Data for "Testing the evolutionary potential of an alpine plant: Phenotypic plasticity in response to growth temperature far outweighs parental environmental effects and other genetic causes of variation"

### Supplementary Data Contents

*Table S1:* Summary statistics for all phenotypic traits.

*Table S2:* Linear regression model output testing fitness index suitability.

*Table S3:* Model outputs for linear selection by growth treatment 1.

*Table S4:* Model outputs for linear selection by growth treatment 2.

*Table S5:* Model outputs for quadratic selection by growth treatment 1.

*Table S6:* Model outputs for quadratic selection by growth treatment 2.

*Fig. S1:* Summary of the multi-generational breeding design.

*Fig. S2:* Correlation between total reproductive stems and reproductive mass.

*Fig. S3:* Reaction norms of phenotypic traits.

*Fig. S4:* Isolated parental temperature effects for phenotypic traits.

*Fig. S5:* Matched/mismatched parent-offspring environment effects on traits 1.

*Fig. S6:* Matched/mismatched parent-offspring environment effects on traits 2.

*Fig. S7:* Tests for inbreeding depression in phenotypic traits 1.

*Fig. S8:* Tests for inbreeding depression in phenotypic traits 2.

**Table S1:** Summary statistics for all phenotypic traits across each treatment combination.

| Phenotypic trait | Growth temperature | Parental temperature | Outcross/ Self | *n* | Mean | S.E.M. |
|---|---|---|---|---|---|---|
| **Total reproductive stems (*n*)** | **Overall** | **Overall** | **Overall** | **1023** | **37.37** | **0.72** |
| | Cool | Cool | Outcross | 184 | 36.16 | 1.78 |
| | Cool | Cool | Self | 90 | 28.93 | 2.29 |
| | Cool | Warm | Outcross | 171 | 35.89 | 2.01 |
| | Cool | Warm | Self | 66 | 32.83 | 2.51 |
| | Warm | Cool | Outcross | 185 | 44.17 | 1.84 |
| | Warm | Cool | Self | 92 | 34.15 | 2.28 |
| | Warm | Warm | Outcross | 171 | 42.49 | 1.87 |
| | Warm | Warm | Self | 64 | 32.56 | 2.90 |
| **Flowering onset (days)** | **Overall** | **Overall** | **Overall** | **1003** | **41.93** | **0.23** |
| | Cool | Cool | Outcross | 182 | 45.34 | 0.43 |
| | Cool | Cool | Self | 85 | 46.54 | 0.77 |
| | Cool | Warm | Outcross | 167 | 46.28 | 0.49 |
| | Cool | Warm | Self | 63 | 48.38 | 0.84 |
| | Warm | Cool | Outcross | 184 | 36.80 | 0.41 |
| | Warm | Cool | Self | 91 | 38.41 | 0.74 |
| | Warm | Warm | Outcross | 168 | 36.91 | 0.37 |
| | Warm | Warm | Self | 63 | 41.33 | 1.17 |
| **Dry above-ground biomass (g)** | **Overall** | **Overall** | **Overall** | **988** | **9.82** | **0.11** |
| | Cool | Cool | Outcross | 178 | 9.52 | 0.26 |
| | Cool | Cool | Self | 86 | 8.45 | 0.39 |
| | Cool | Warm | Outcross | 167 | 9.64 | 0.28 |
| | Cool | Warm | Self | 64 | 8.86 | 0.41 |
| | Warm | Cool | Outcross | 181 | 10.80 | 0.31 |
| | Warm | Cool | Self | 85 | 9.15 | 0.45 |
| | Warm | Warm | Outcross | 165 | 10.90 | 0.32 |
| | Warm | Warm | Self | 62 | 9.12 | 0.49 |
| **Chl. content (SPAD units)** | **Overall** | **Overall** | **Overall** | **984** | **35.44** | **0.19** |
| | Cool | Cool | Outcross | 178 | 33.36 | 0.44 |
| | Cool | Cool | Self | 86 | 34.81 | 0.65 |
| | Cool | Warm | Outcross | 167 | 33.39 | 0.46 |
| | Cool | Warm | Self | 62 | 34.14 | 0.61 |
| | Warm | Cool | Outcross | 181 | 36.93 | 0.47 |
| | Warm | Cool | Self | 84 | 37.68 | 0.74 |
| | Warm | Warm | Outcross | 165 | 36.86 | 0.50 |
| | Warm | Warm | Self | 61 | 37.99 | 0.80 |
| **LMA (mg $cm^{-2}$)** | **Overall** | **Overall** | **Overall** | **975** | **1.17** | **0.01** |
| | Cool | Cool | Outcross | 176 | 1.11 | 0.02 |
| | Cool | Cool | Self | 84 | 1.12 | 0.03 |
| | Cool | Warm | Outcross | 166 | 1.11 | 0.02 |
| | Cool | Warm | Self | 62 | 1.12 | 0.03 |
| | Warm | Cool | Outcross | 179 | 1.24 | 0.03 |
| | Warm | Cool | Self | 84 | 1.25 | 0.04 |
| | Warm | Warm | Outcross | 164 | 1.20 | 0.02 |
| | Warm | Warm | Self | 60 | 1.21 | 0.05 |

| Phenotypic trait | Growth temperature | Parental temperature | Outcross/ Self | *n* | Mean | S.E.M. |
|---|---|---|---|---|---|---|
| $F_V/F_M$ | **Overall** | **Overall** | **Overall** | **717** | **0.73** | **0.001** |
| | Cool | Cool | Outcross | 120 | 0.73 | 0.003 |
| | Cool | Cool | Self | 58 | 0.73 | 0.005 |
| | Cool | Warm | Outcross | 117 | 0.73 | 0.003 |
| | Cool | Warm | Self | 45 | 0.73 | 0.006 |
| | Warm | Cool | Outcross | 136 | 0.74 | 0.004 |
| | Warm | Cool | Self | 64 | 0.74 | 0.005 |
| | Warm | Warm | Outcross | 128 | 0.74 | 0.003 |
| | Warm | Warm | Self | 49 | 0.74 | 0.005 |
| **ϕPSII** | **Overall** | **Overall** | **Overall** | **717** | **0.28** | **0.003** |
| | Cool | Cool | Outcross | 120 | 0.25 | 0.010 |
| | Cool | Cool | Self | 58 | 0.27 | 0.014 |
| | Cool | Warm | Outcross | 117 | 0.26 | 0.010 |
| | Cool | Warm | Self | 45 | 0.27 | 0.017 |
| | Warm | Cool | Outcross | 136 | 0.30 | 0.009 |
| | Warm | Cool | Self | 64 | 0.29 | 0.012 |
| | Warm | Warm | Outcross | 128 | 0.29 | 0.009 |
| | Warm | Warm | Self | 49 | 0.31 | 0.014 |
| $T_{crit\text{-}hot}$ **(°C)** | **Overall** | **Overall** | **Overall** | **685** | **44.68** | **0.05** |
| | Cool | Cool | Outcross | 110 | 44.56 | 0.15 |
| | Cool | Cool | Self | 55 | 44.51 | 0.25 |
| | Cool | Warm | Outcross | 108 | 44.46 | 0.15 |
| | Cool | Warm | Self | 44 | 44.72 | 0.31 |
| | Warm | Cool | Outcross | 132 | 44.79 | 0.17 |
| | Warm | Cool | Self | 64 | 44.76 | 0.26 |
| | Warm | Warm | Outcross | 124 | 44.85 | 0.17 |
| | Warm | Warm | Self | 48 | 44.77 | 0.31 |
| $T_{crit\text{-}cold}$ **(°C)** | **Overall** | **Overall** | **Overall** | **707** | **-13.12** | **0.07** |
| | Cool | Cool | Outcross | 116 | -13.46 | 0.19 |
| | Cool | Cool | Self | 57 | -13.46 | 0.33 |
| | Cool | Warm | Outcross | 116 | -13.20 | 0.21 |
| | Cool | Warm | Self | 45 | -13.42 | 0.30 |
| | Warm | Cool | Outcross | 136 | -13.02 | 0.22 |
| | Warm | Cool | Self | 61 | -12.67 | 0.31 |
| | Warm | Warm | Outcross | 128 | -12.65 | 0.22 |
| | Warm | Warm | Self | 48 | -13.49 | 0.31 |

**Table S2:** Linear regression model output testing the suitability of total number of reproductive stems as a fitness index. Reproductive mass = harvested capsule mass × number of capsules weighed. See also Fig. S2.

| **Response: Total number of reproductive stems** | | | |
|---|---|---|---|
| ***Predictor*** | ***Estimate*** | ***95% CI*** | ***P*** |
| (Intercept) | 13.53 | 6.42 – 20.64 | **<0.001** |
| Reproductive mass | 84.83 | 71.09 – 98.57 | **<0.001** |
| Growth temperature (warm) | -3.65 | -13.04 – 5.75 | 0.443 |
| Parent temperature (warm) | -6.56 | -18.24 – 5.12 | 0.267 |
| Cross type (self) | 2.39 | -2.67 – 7.45 | 0.350 |
| Reproductive mass × Growth temperature (warm) | 20.82 | -3.19 – 44.82 | 0.088 |
| Reproductive mass × Parent temperature (warm) | 4.59 | -21.70 – 30.89 | 0.729 |
| Growth temperature (warm) × Parent temperature (warm) | 1.67 | -15.58 – 18.92 | 0.848 |
| (Reproductive mass × Growth temperature (warm)) × Parent temperature (warm) | 30.16 | -15.94 – 76.26 | 0.197 |
| Observations (*n*) | 100 | | |
| $R^2$ | 0.822 | | |

**Table S3:** Full model output for testing whether linear selection (β) differs by growth treatment for four phenotypic traits. Significant interaction terms indicates that selection varies depending on growth temperature treatment (i.e., that selection on cool-grown and warm-grown plants differs for that trait).

| ***Response variable: w*** | *** Flowering onset (β)** | *** Biomass (β)** | *** Chlorophyll content (β)** | *** LMA (β)** |
|---|---|---|---|---|
| ***Fixed effects*** | | | | |
| ***Estimate [95% CI]*** | | | | |
| Intercept (cool Growth temp., cool Parental temp., outcrossed) | **1.092 [0.903, 1.290]** | **1.188 [1.030, 1.340]** | **0.959 [0.762, 1.146]** | **0.980 [0.781, 1.172]** |
| * Trait (β) | **-0.215 [-0.270, -0.162]** | **0.451 [0.400, 0.503]** | -0.017 [-0.061, 0.026] | **0.055 [0.011, 0.099]** |
| Growth temp. (warm) | **-0.149 [-0.223, -0.076]** | 0.031 [-0.016, 0.079] | **0.094 [0.033, 0.156]** | **0.080 [0.022, 0.140]** |
| Parental temp. (warm) | 0.020 [-0.063, 0.102] | -0.055 [-0.148, 0.035] | -0.044 [-0.137, 0.047] | -0.036 [-0.133, 0.059] |
| Cross type (self-pollinated) | **-0.088 [-0.169, -0.009]** | -0.059 [-0.137, 0.017] | **-0.189 [-0.273, -0.109]** | **-0.194 [-0.278, -0.114]** |
| Harvest date | -- | **-0.248 [-0.316, -0.181]** | **0.187 [0.124, 0.253]** | 0.165 [0.101, 0.231] |
| Growth temp. (warm) × trait (β) | ***-0.097 [-0.171, -0.023]*** | ***-0.093 [-0.144, -0.044]*** | 0.011 [-0.047, 0.069] | -0.037 [-0.092, 0.018] |
| ***Random effects: variance components*** | | | | |
| ***Estimate (SD) [95% CI]*** | | | | |
| $V_B$ intercept (block) | 0.128 [0.029, 0.468] | 0.098 [0.019, 0.377] | 0.137 [0.034, 0.480] | 0.136 [0.030, 0.504] |
| $V_A$ intercept (additive genetic) | 0.129 [0.065, 0.183] | 0.155 [0.095, 0.210] | 0.121 [0.022, 0.197] | 0.122 [0.021, 0.203] |
| $V_M$ intercept (maternal) | 0.052 [0.002, 0.126] | 0.111 [0.013, 0.186] | 0.115 [0.016, 0.193] | 0.116 [0.014, 0.191] |
| $V_R$ (residual) | 0.537 [0.510, 0.566] | 0.434 [0.411, 0.457] | 0.572 [0.542, 0.603] | 0.566 [0.536, 0.597] |

Values in bold represent fixed effects that have 95% CIs that are distinct from zero. The models include interaction terms between trait × growth temperature treatment for β to test whether selection differs by growth treatment. Italics indicates that the respective interaction term has 95% CIs that are distinct from zero, which indicates that selection varies depending on growth temperature treatment (i.e., that selection on cool-grown and warm-grown plants differs for that trait).

**Table S4:** Full model output for testing whether linear selection (β) differs by growth treatment for four phenotypic traits. Significant interaction terms indicates that selection varies depending on growth temperature treatment (i.e., that selection on cool-grown and warm-grown plants differs for that trait).

| ***Response variable: w*** | *** $F_V/F_M$ (β)** | *** ϕPSII (β)** | *** $T_{crit\text{-}hot}$ (β)** | *** $T_{crit\text{-}cold}$ (β)** |
|---|---|---|---|---|
| ***Fixed effects*** | | | | |
| ***Estimate [95% CI]*** | | | | |
| Intercept (cool Growth temp., cool Parental temp., outcrossed) | **0.997 [0.790, 1.205]** | **0.996 [0.806, 1.175]** | **0.991 [0.810, 1.172]** | **0.981 [0.799, 1.150]** |
| * Trait (β) | 0.023 [-0.027, 0.075] | 0.026 [-0.020, 0.073] | -0.030 [-0.083, 0.024] | -0.056 [-0.111, 0.002] |
| Growth temp. (warm) | 0.053 [-0.013, 0.118] | 0.049 [-0.017, 0.114] | 0.065 [0.000, 0.133] | 0.061 [-0.004, 0.127] |
| Parental temp. (warm) | -0.052 [-0.152, 0.046] | -0.053 [-0.155, 0.049] | -0.050 [-0.148, 0.051] | -0.048 [-0.149, 0.052] |
| Cross type (self-pollinated) | **-0.150 [-0.240, -0.062]** | **-0.151 [-0.245, -0.060]** | **-0.165 [-0.254, -0.078]** | **-0.143 [-0.233, -0.055]** |
| Growth temp. (warm) × trait (β) | -0.032 [-0.103, 0.038] | -0.021 [-0.086, 0.043] | 0.001 [-0.068, 0.071] | 0.051 [-0.019, 0.120] |
| Hail (measured post-hail) | **0.189 [0.056, 0.308]** | **0.193 [0.068, 0.311]** | **0.247 [0.126, 0.362]** | **0.186 [0.067, 0.303]** |
| ***Random effects: variance components*** | | | | |
| ***Estimate (SD) [95% CI]*** | | | | |
| $V_B$ intercept (block) | 0.111 [0.002, 0.712] | 0.089 [0.002, 0.531] | 0.087 [0.001, 0.517] | 0.091 [0.002, 0.517] |
| $V_A$ intercept (additive genetic) | 0.140 [0.046, 0.214] | 0.140 [0.040, 0.215] | 0.133 [0.035, 0.211] | 0.151 [0.055, 0.218] |
| $V_M$ intercept (maternal) | 0.099 [0.006, 0.189] | 0.100 [0.007, 0.192] | 0.104 [0.008, 0.196] | 0.089 [0.004, 0.181] |
| $V_R$ (residual) | 0.579 [0.544, 0.617] | 0.578 [0.543, 0.615] | 0.576 [0.542, 0.614] | 0.577 [0.543, 0.614] |

Values in bold represent fixed effects that have 95% CIs that are distinct from zero. The models include interaction terms between trait × growth temperature treatment for β to test whether selection differs by growth treatment. Italics indicates that the respective interaction term has 95% CIs that are distinct from zero, which indicates that selection varies depending on growth temperature treatment (i.e., that selection on cool-grown and warm-grown plants differs for that trait).

**Table S5:** Full model output for testing whether quadratic selection (γ) differs by growth treatment for four phenotypic traits. Significant interaction terms indicates that selection varies depending on growth temperature treatment (i.e., that selection on cool-grown and warm-grown plants differs for that trait).

| ***Response variable: w*** | *** Flowering onset (γ)** | *** Biomass (γ)** | *** Chlorophyll content (γ)** | *** LMA (γ)** |
|---|---|---|---|---|
| ***Fixed effects*** | | | | |
| ***Estimate [95% CI]*** | | | | |
| Intercept (cool Growth temp., cool Parental temp., outcrossed) | **1.084 [0.899, 1.261]** | **1.189 [1.038, 1.333]** | **0.944 [0.734, 1.148]** | **0.977 [0.774, 1.159]** |
| Trait | -0.204 [-0.606, 0.234] | **0.356 [0.190, 0.529]** | 0.244 [-0.127, 0.634] | **0.306 [0.054, 0.588]** |
| * Trait$^2$ (γ) | -0.012 [-0.433, 0.366] | 0.121 [-0.065, 0.300] | -0.281 [-0.696, 0.114] | -0.259 [-0.559, 0.000] |
| Growth temp. (warm) | **-0.143 [-0.236, -0.053]** | 0.062 [0.000, 0.124] | **0.108 [0.028, 0.190]** | **0.100 [0.018, 0.181]** |
| Parental temp. (warm) | 0.028 [-0.069, 0.126] | -0.039 [-0.145, 0.067] | -0.037 [-0.145, 0.073] | -0.035 [-0.150, 0.076] |
| Cross type (self-pollinated) | **-0.090 [-0.166, -0.014]** | **-0.048 [-0.128, 0.032]** | **-0.188 [-0.270, -0.106]** | **-0.188 [-0.276, -0.104]** |
| Harvest date | -- | **-0.274 [-0.344, -0.205]** | **0.194 [0.130, 0.259]** | **0.149 [0.081, 0.217]** |
| Growth temp. (warm) × trait | **-0.606 [-1.137, -0.102]** | 0.315 [0.098, 0.524] | -0.046 [-0.562, 0.480] | 0.093 [-0.286, 0.455] |
| Growth temp. (warm) × trait$^2$ (γ) | ***0.541 [0.054, 1.054]*** | ***-0.410 [-0.630, -0.183]*** | 0.083 [-0.450, 0.610] | -0.112 [-0.472, 0.273] |
| Growth temp. (warm) × parental temp. (warm) | **-0.015 [-0.124, 0.097]** | -0.048 [-0.137, 0.043] | -0.014 [-0.128, 0.099] | -0.017 [-0.130, 0.100] |
| ***Random effects: variance components*** | | | | |
| ***Estimate (SD) [95% CI]*** | | | | |
| $V_B$ intercept (block) | 0.127 [0.028, 0.470] | 0.093 [0.016, 0.343] | 0.146 [0.034, 0.539] | 0.130 [0.028, 0.465] |
| $V_A$ intercept (additive genetic) | 0.124 [0.067, 0.177] | 0.160 [0.097, 0.218] | 0.114 [0.020, 0.189] | 0.138 [0.043, 0.213] |
| $V_M$ intercept (maternal) | 0.050 [0.003, 0.120] | 0.117 [0.016, 0.196] | 0.117 [0.013, 0.193] | 0.110 [0.011, 0.193] |
| $V_R$ (residual) | 0.535 [0.508. 0.564] | 0.427 [0.405, 0.450] | 0.572 [0.542, 0.603] | 0.558 [0.528, 0.589] |

Values in bold represent fixed effects that have 95% CIs that are distinct from zero. The models include interaction terms between trait × growth temperature treatment for β and trait$^2$ × growth temperature treatment for γ to test whether selection differs by growth treatment. Italics indicates that the respective interaction term has 95% CIs that are distinct from zero, which indicates that selection varies depending on growth temperature treatment (i.e., that selection on cool-grown and warm-grown plants differs for that trait).

**Table S6:** Full model output for testing whether quadratic selection (γ) differs by growth treatment for four phenotypic traits. Significant interaction terms indicates that selection varies depending on growth temperature treatment (i.e., that selection on cool-grown and warm-grown plants differs for that trait).

| ***Response variable: w*** | *** $F_V/F_M$ (γ)** | *** ϕPSII (γ)** | *** $T_{crit\text{-}hot}$ (γ)** | *** $T_{crit\text{-}cold}$ (γ)** |
|---|---|---|---|---|
| ***Fixed effects*** | | | | |
| ***Estimate [95% CI]*** | | | | |
| Intercept (cool Growth temp., cool Parental temp., outcrossed) | **0.978 [0.777, 1.170]** | **0.981 [0.784, 1.185]** | **0.988 [0.778, 1.211]** | **0.960 [0.776, 1.136]** |
| Trait (β) | 0.221 [-1.039, 1.804] | 0.079 [-0.122, 0.294] | -0.583 [-2.337, 1.319] | 0.142 [-0.259, 0.517] |
| * Trait$^2$ (γ) | -0.197 [-1.764, 1.065] | -0.056 [-0.283, 0.159] | 0.553 [-1.343, 2.306] | 0.200 [-0.193, 0.567] |
| Growth temp. (warm) | 0.081 [-0.007, 0.172] | 0.073 [-0.015, 0.165] | 0.078 [-0.012, 0.167] | **0.100 [0.010, 0.189]** |
| Parental temp. (warm) | -0.024 [-0.142, 0.097] | -0.025 [-0.151, 0.095] | -0.037 [-0.159, 0.081] | -0.007 [-0.130, 0.112] |
| Cross type (self-pollinated) | **-0.147 [-0.242, -0.057]** | **-0.142 [-0.237, -0.052]** | **-0.168 [-0.256, -0.081]** | **-0.141 [-0.234, -0.053]** |
| Hail (measured post-hail) | **0.192 [0.060, 0.312]** | **0.205 [0.082, 0.322]** | **0.244 [0.122, 0.364]** | **0.186 [0.064, 0.302]** |
| Growth temp. (warm) × trait | -0.733 [-2.613, 0.948] | 0.176 [-0.158, 0.505] | 0.119 [-2.069, 2.254] | -0.376 [-0.888, 0.133] |
| Growth temp. (warm) × trait$^2$ (γ) | 0.697 [-0.966, 2.560] | -0.181 [-0.508, 0.146] | -0.118 [-2.255, 2.061] | -0.433 [-0.941, 0.074] |
| Growth temp. (warm) × parental temp. (warm) | -0.059 [-0.185, 0.071] | -0.064 [-0.195, 0.066] | -0.028 [-0.152, 0.096] | -0.079 [-0.204, 0.050] |
| ***Random effects: variance components*** | | | | |
| ***Estimate (SD) [95% CI]*** | | | | |
| $V_B$ intercept (block) | 0.100 [0.001, 0.588] | 0.100 [0.001, 0.617] | 0.109 [0.002, 0.726] | 0.090 [0.002, 0.511] |
| $V_A$ intercept (additive genetic) | 0.132 [0.025, 0.211] | 0.138 [0.033, 0.215] | 0.129 [0.031, 0.208] | 0.158 [0.072, 0.226] |
| $V_M$ intercept (maternal) | 0.106 [0.008, 0.194] | 0.101 [0.008, 0.190] | 0.106 [0.010, 0.194] | 0.089 [0.006, 0.185] |
| $V_R$ (residual) | 0.580 [0.545, 0.617] | 0.577 [0.543, 0.614] | 0.578 [0.542, 0.617] | 0.574 [0.539, 0.612] |

Values in bold represent fixed effects that have 95% CIs that are distinct from zero. The models include interaction terms between trait × growth temperature treatment for β and trait$^2$ × growth temperature treatment for γ to test whether selection differs by growth treatment. Italics indicates that the respective interaction term has 95% CIs that are distinct from zero, which indicates that selection varies depending on growth temperature treatment (i.e., that selection on cool-grown and warm-grown plants differs for that trait).

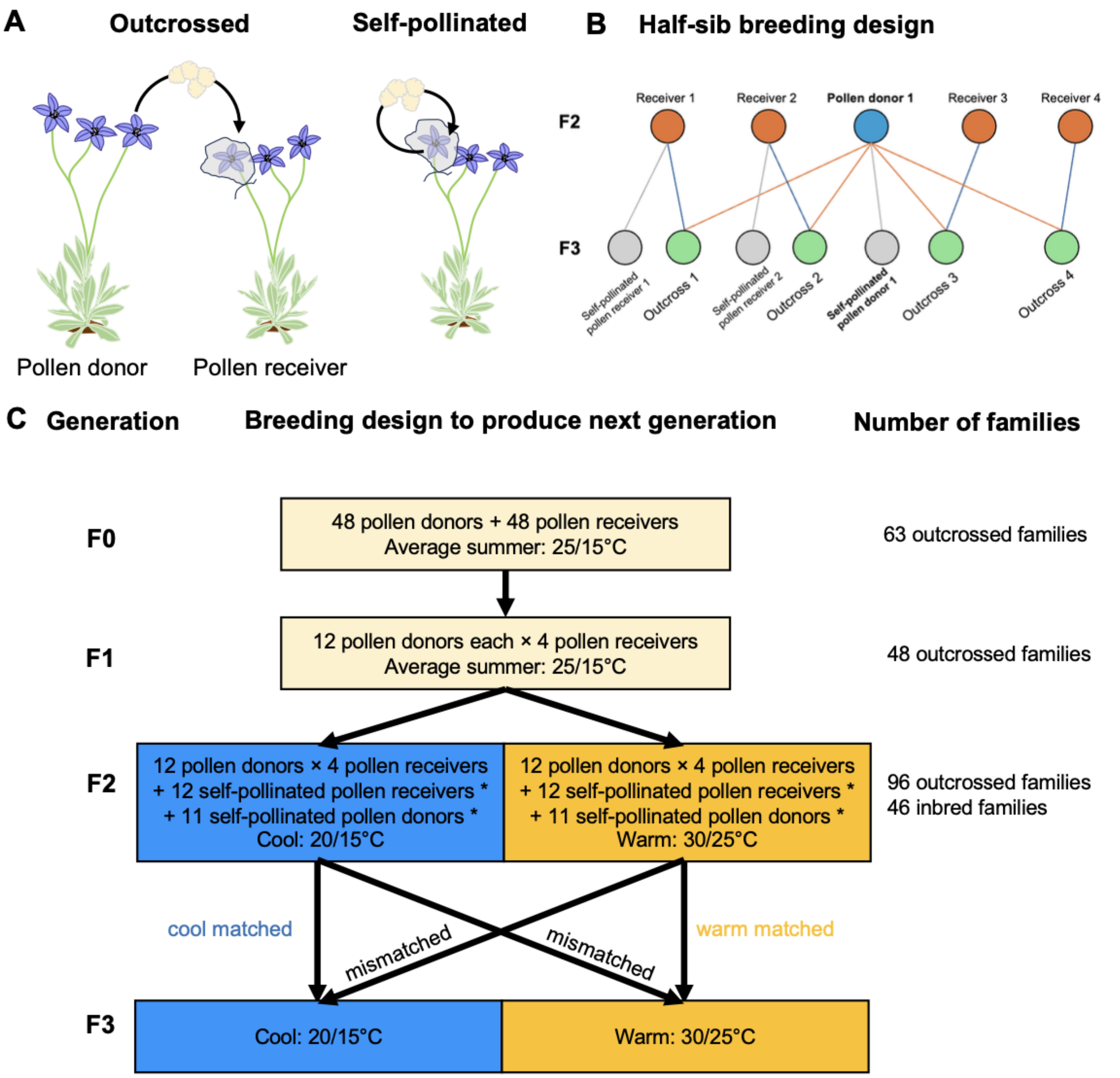


**Fig. S1:** Summary of the multi-generational breeding design to produce the F3 plants that were used in the current study. (A) Hand pollination process for outcrossed families where pollen donor plant donates pollen to the pollen receiver plant, the pollinated flower is bagged, and then the capsule is collected when mature and dry; and for self-pollinated families where pollen is donated and received by the same flower on the same plant. (B) Example of half-sib breeding design where one F2 pollen donor (blue) is crossed with four F2 pollen receivers (orange) to produce F3 outcrossed families (green). Pollen receivers 1 and 2 are also self-pollinated to produce a 'self-pollinated pollen receiver' (grey) and the Pollen donor 1 is self-pollinated to produce a 'self-pollinated pollen donor' (grey, bold text). (C) Visualisation of the breeding design by generation. Rectangles include the breeding design employed to produce the following generation as well as the day/night experimental temperature treatments to which the generation was exposed. The parental and growth temperature treatment of cool and warm are reciprocally matched and mismatched between the F2 (parental) and F3 (growth) generations. Number of families represents the total number of families produced in that given generation.

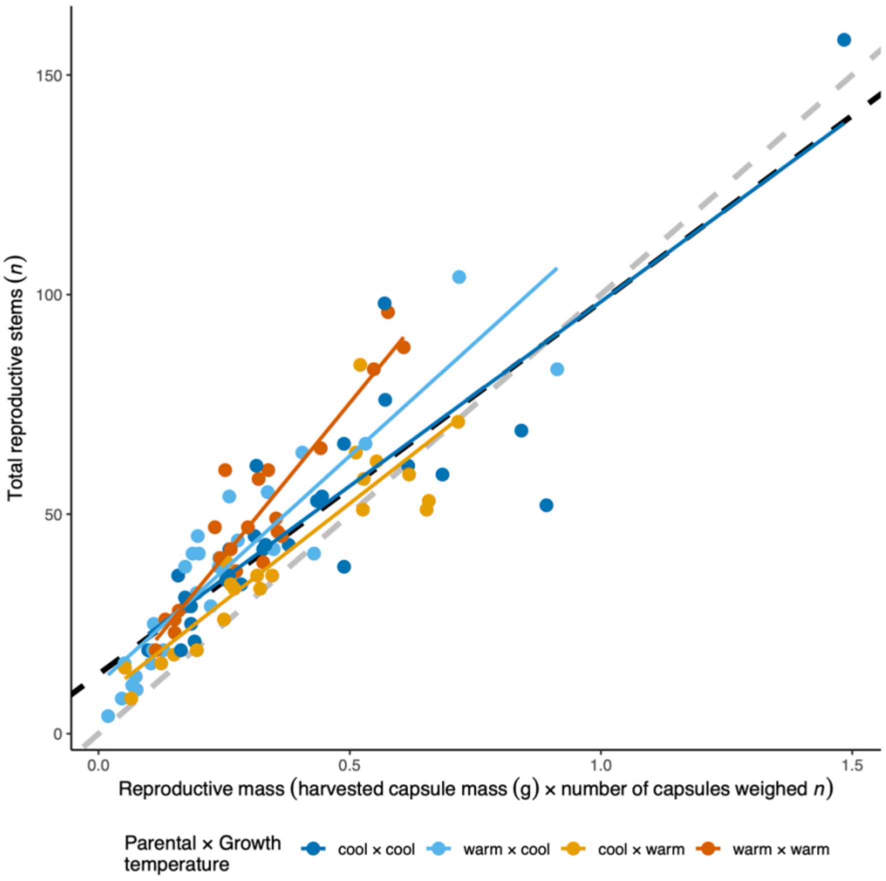


**Fig. S2:** Correlation between the total number of reproductive stems and the harvested capsule mass × number of capsules weighed. The partial regressions of each parental × growth temperature treatment combination are shown by the different colours. Black dashed line represents the overall model fit, and the grey dashed line represents a 1:1 scale.

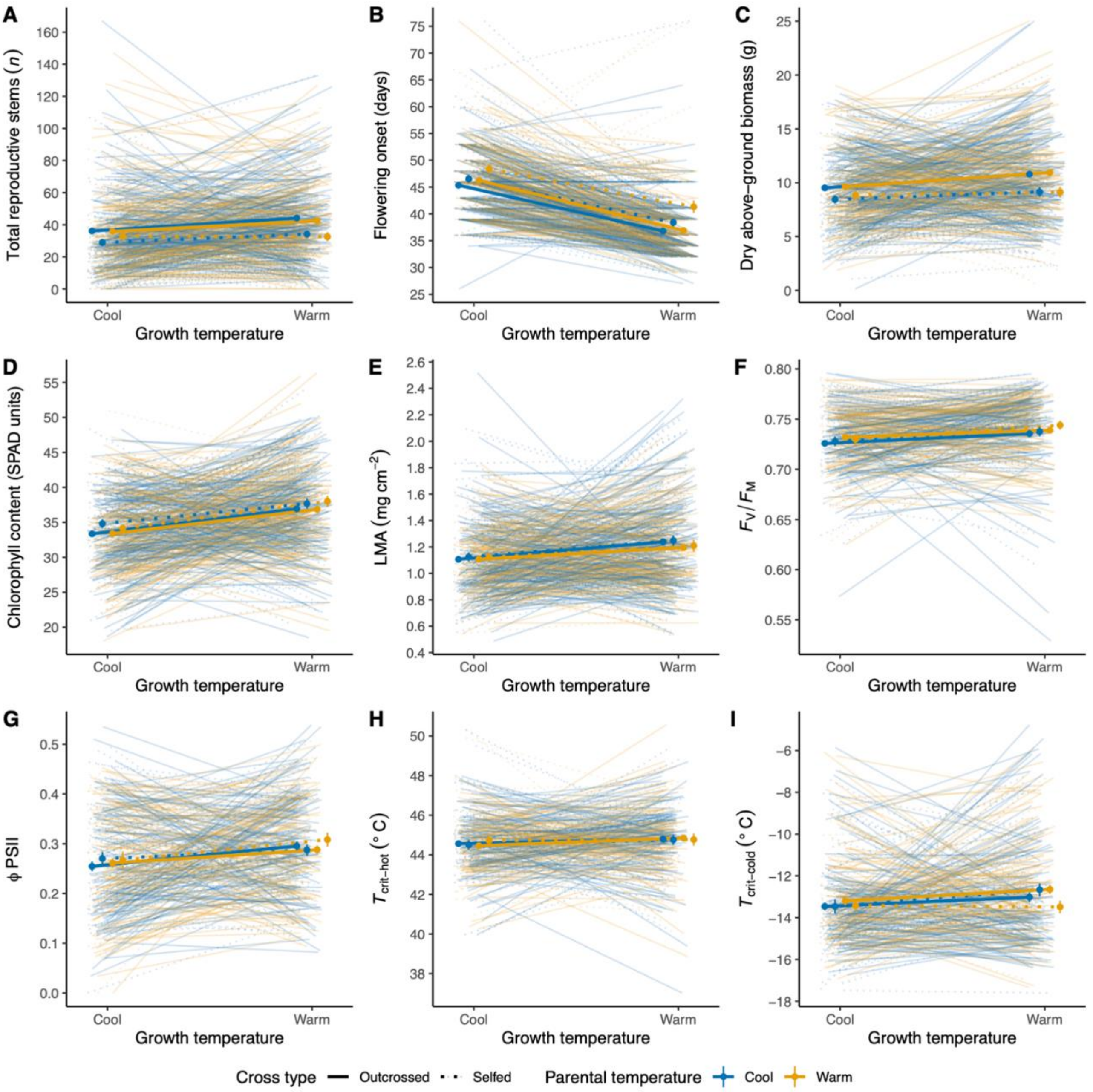


**Fig. S3:** Reaction norms of phenotypic traits: (A) total reproductive stems, (B) flowering onset, (C) biomass, (D) chlorophyll content, (E) LMA, as well as photosystem traits: (F) $F_V/F_M$ and (G) ϕPSII, and thermal tolerance traits: (H) $T_{crit\text{-}hot}$ and (I) $T_{crit\text{-}cold}$ in response to growth temperature treatments. Within each cool and warm growth temperature treatment, plants were grown under an environment that was either cool (blues) or warm (oranges) and were offspring plants were from either outcrossed (solid lines) or self-pollinated (dotted lines) parents. Thicker lines represent the population-level means, while lighter lines represent paired individuals of the same parentage. Points and error bars represent means ± S.E. of the raw data. Model outputs are shown in Tables 1 and 2 and overall population-level means are shown in Fig. 1.

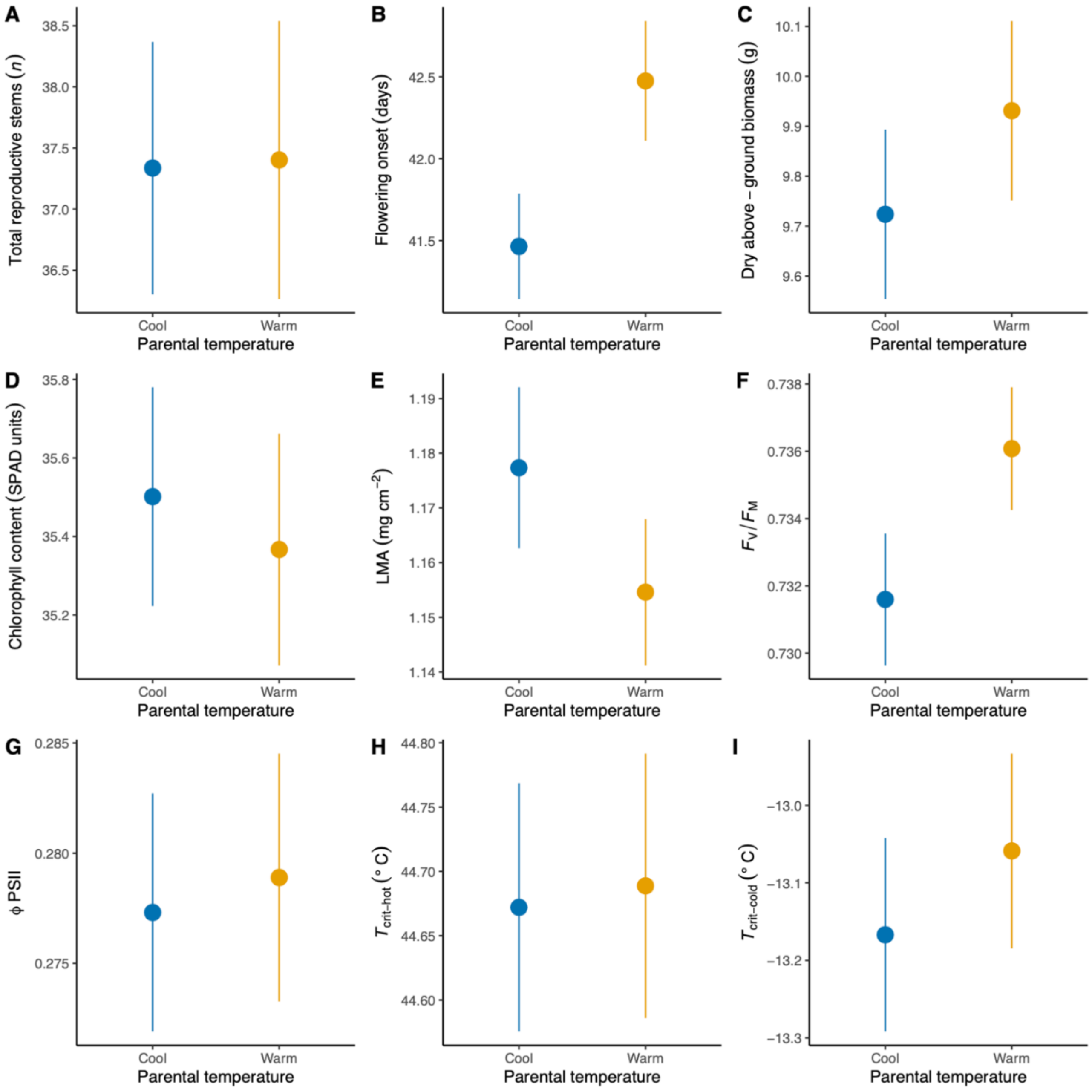


**Fig. S4:** Isolated parental temperature effects for fitness and phenotypic traits: (A) total reproductive stems, (B) flowering onset, (C) biomass, (D) chlorophyll content, (E) LMA, as well as photosystem traits: (F) $F_V/F_M$ and (G) ϕPSII, and thermal tolerance traits: (H) $T_{crit\text{-}hot}$ and (I) $T_{crit\text{-}cold}$. Points and error bars represent means ± S.E. of the raw data.

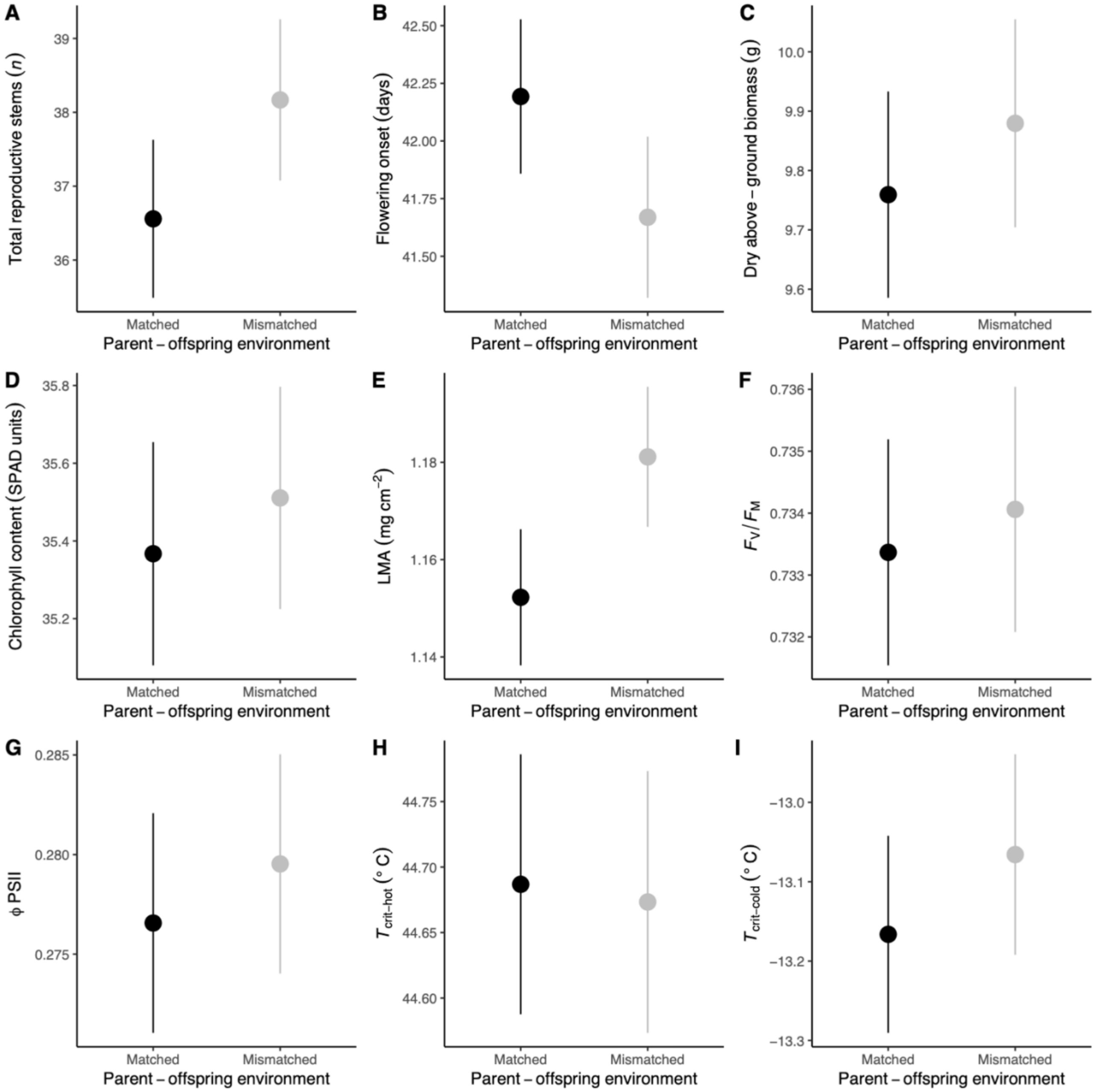


**Fig. S5:** Mean differences between matched and mismatched parent-offspring environments for fitness and phenotypic traits: (A) total reproductive stems, (B) flowering onset, (C) biomass, (D) chlorophyll content, (E) LMA, as well as photosystem traits: (F) $F_V/F_M$ and (G) ϕPSII, and thermal tolerance traits: (H) $T_{crit\text{-}hot}$ and (I) $T_{crit\text{-}cold}$. Matched refers to when growing conditions of both the parent and offspring (growth for the plants being phenotyped) are the same (i.e., parental cool × growth cool and parental warm × growth warm) and mismatch refers to when they are different (i.e., parental warm × growth cool and parental cool × growth warm). Points and error bars represent means ± S.E. of the raw data.

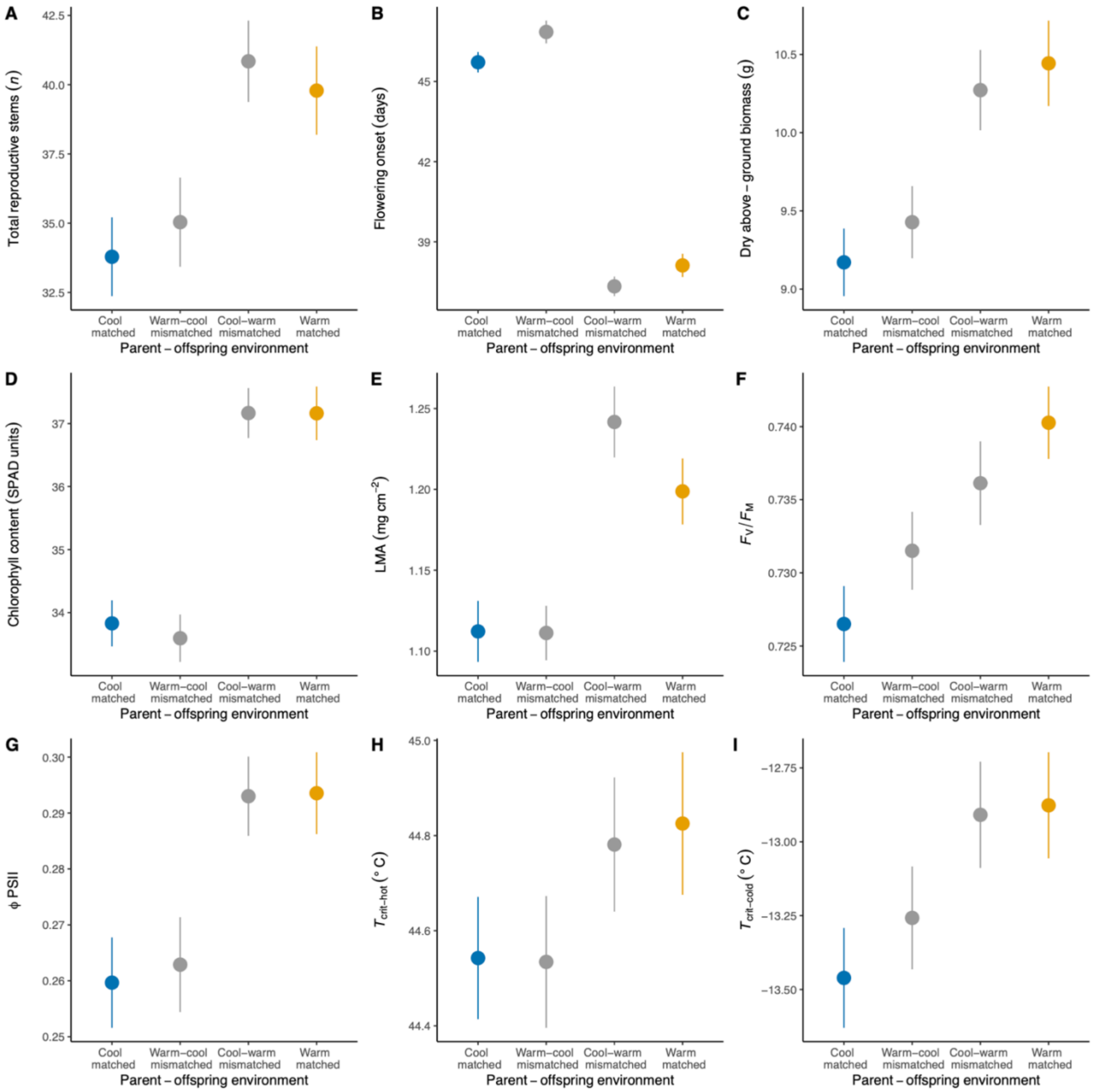


**Fig. S6:** Mean differences between each specific matched and mismatched parent-offspring environments for fitness and phenotypic traits: (A) total reproductive stems, (B) flowering onset, (C) biomass, (D) chlorophyll content, (E) LMA, as well as photosystem traits: (F) $F_V/F_M$ and (G) ϕPSII, and thermal tolerance traits: (H) $T_{crit\text{-}hot}$ and (I) $T_{crit\text{-}cold}$. Matched refers to when growing conditions of both the parent and offspring (growth for the plants being phenotyped) are the same (i.e., parental cool × growth cool and parental warm × growth warm) and mismatch refers to when they are different (i.e., parental warm × growth cool and parental cool × growth warm). Points and error bars represent means ± S.E. of the raw data.

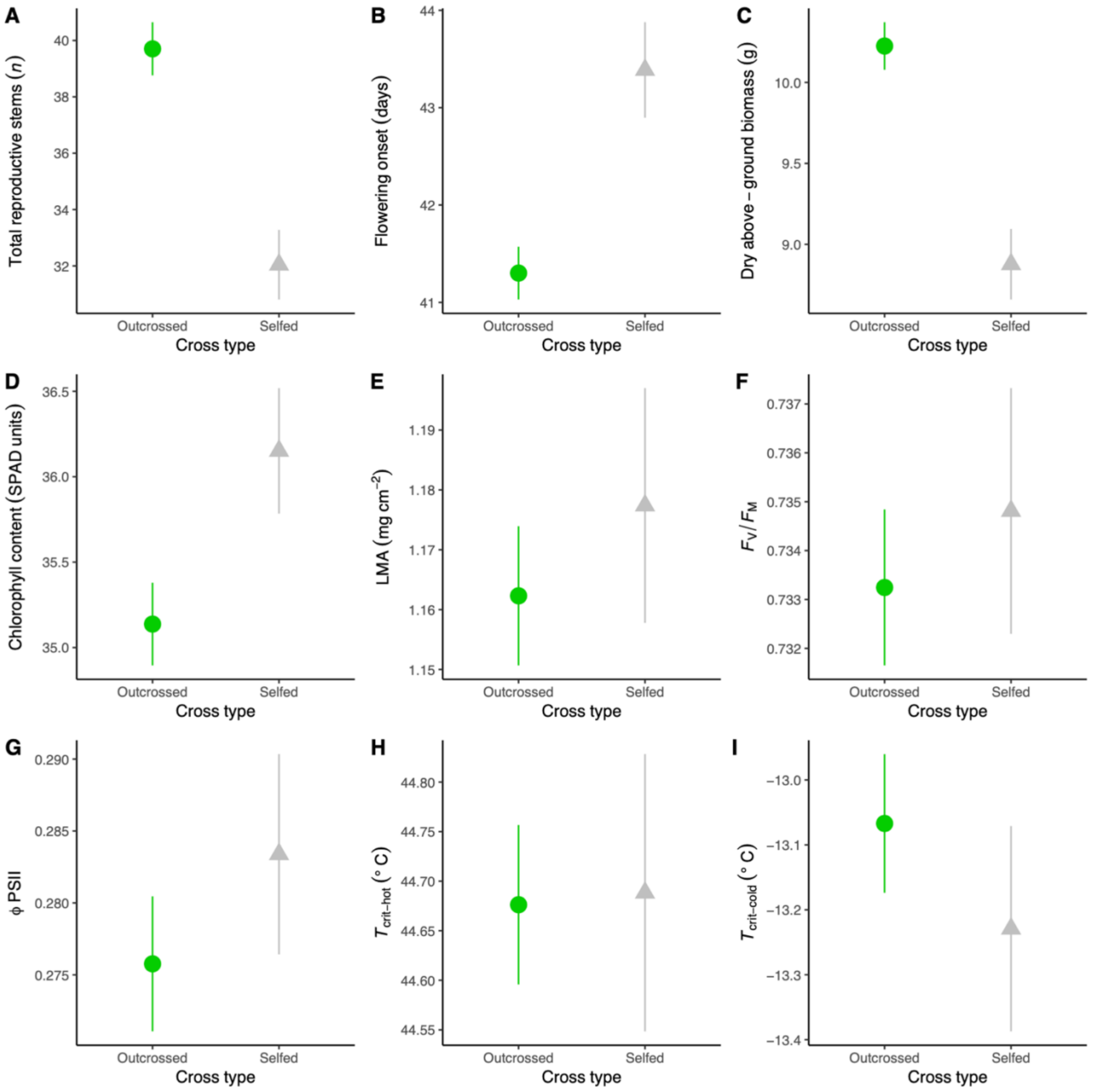


**Fig. S7:** Tests for inbreeding depression. Mean differences between outcrossed and self-pollinated plants overall: (A) total reproductive stems, (B) flowering onset, (C) biomass, (D) chlorophyll content, (E) LMA, as well as photosystem traits: (F) $F_V/F_M$ and (G) ϕPSII, and thermal tolerance traits: (H) $T_{crit\text{-}hot}$ and (I) $T_{crit\text{-}cold}$. Points and error bars represent means ± S.E. of the raw data. Green circles represent plants produced from outcrosses and grey triangles represent plants produced from self-pollination.

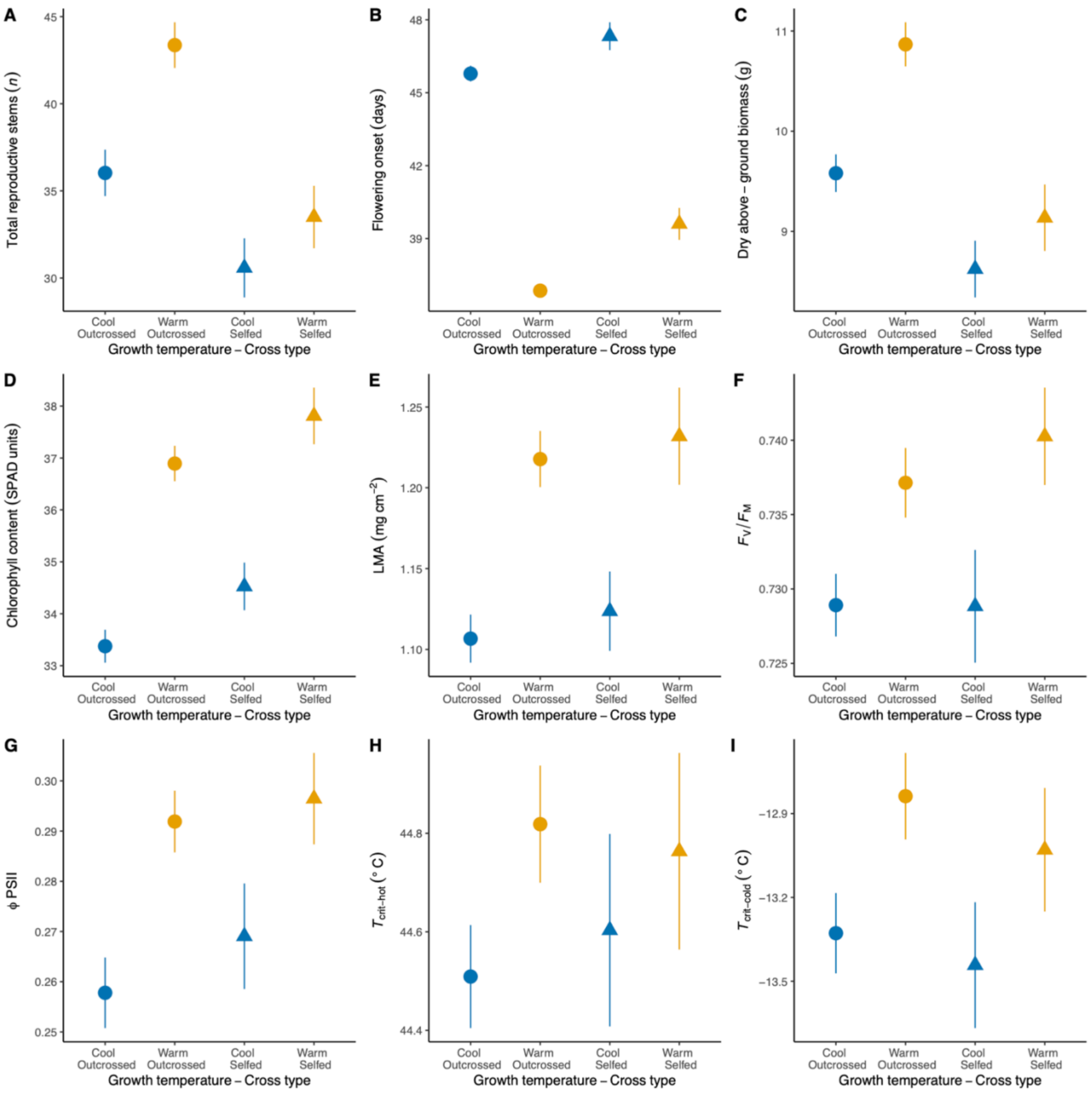


**Fig. S8:** Tests for inbreeding depression. Mean differences between outcrossed and self-pollinated plants under cool (blue) and warm (orange) growth temperatures for fitness and phenotypic traits: (A) total reproductive stems, (B) flowering onset, (C) biomass, (D) chlorophyll content, (E) LMA, as well as photosystem traits: (F) $F_V/F_M$ and (G) ϕPSII, and thermal tolerance traits: (H) $T_{crit\text{-}hot}$ and (I) $T_{crit\text{-}cold}$. Points and error bars represent means ± S.E. of the raw data. Circles represent plants produced from outcrosses and triangles represent plants produced from self-pollination. There were no interactions between growth temperature and cross type that had credible intervals distinct from zero in full models for any trait. Therefore, the effects of inbreeding depression are relatively consistent in both growth temperatures, i.e., the magnitude of change in trait value due to inbreeding is similar in both cool and warm growth temperatures.
